# A multidrug-resistant *Salmonella enterica* Typhimurium DT104 complex lineage circulating among humans and cattle in the United States lost the ability to produce pertussis-like toxin ArtAB

**DOI:** 10.1101/2022.04.06.487395

**Authors:** Laura M. Carroll, Nicolo Piacenza, Rachel A. Cheng, Martin Wiedmann, Claudia Guldimann

## Abstract

*Salmonella enterica* subspecies *enterica* serotype Typhimurium definitive type 104 (DT104) can infect both humans and animals and is often multidrug-resistant (MDR). Previous studies have indicated that, unlike most *S.* Typhimurium, the overwhelming majority of DT104 strains produce pertussis-like toxin ArtAB via prophage-encoded genes *artAB*. However, DT104 that lack *artAB* have been described on occasion. Here, we identify a MDR DT104 complex lineage circulating among humans and cattle in the United States, which lacks *artAB* (i.e., the “U.S. *artAB*-negative major clade”; *n* = 42 genomes). Unlike most other bovine- and human-associated DT104 complex strains from the U.S. (*n* = 230 total genomes), which harbor *artAB* on prophage Gifsy-1 (*n* = 177), members of the U.S. *artAB*-negative major clade lack Gifsy-1, as well as anti-inflammatory effector *gogB*. The U.S. *artAB*-negative major clade encompasses human- and cattle-associated strains isolated from ≥11 U.S. states over a twenty-year period. The clade was predicted to have lost *artAB*, Gifsy-1, and *gogB* circa 1985-1987 (95% highest posterior density interval 1979.0-1992.1). When compared to DT104 genomes from other world regions (*n* = 752 total genomes), several additional, sporadic *artAB*, Gifsy-1, and/or *gogB* loss events among clades encompassing ≤5 genomes were observed. Using phenotypic assays that simulate conditions encountered during human and/or bovine digestion, members of the U.S. *artAB*-negative major clade did not differ from closely related Gifsy-1/*artAB*/*gogB*-harboring U.S. DT104 complex strains (ANOVA raw *P*-value > 0.05); thus, future research is needed to elucidate the roles that *artAB*, *gogB*, and Gifsy-1 play in DT104 virulence in humans and animals.

**Impact Statement:** Multi-drug resistant (MDR) *Salmonella enterica* serotype Typhimurium definitive type 104 (DT104) was responsible for a global epidemic among humans and animals throughout the 1990s and continues to circulate worldwide. Previous studies have indicated that the vast majority of DT104 produce pertussis-like toxin ArtAB via prophage-encoded *artAB*. Here, we identify a DT104 complex lineage that has been circulating among cattle and humans across ≥11 U.S. states for over twenty years, which lacks the ability to produce ArtAB (i.e., the “U.S. *artAB*-negative major clade”). The common ancestor of all U.S. *artAB*-negative major clade members lost the ability to produce ArtAB in the 1980s; however, the reason for this loss-of-function event within this well-established pathogen remains unclear. The role that ArtAB plays in DT104 virulence remains elusive, and phenotypic assays conducted here indicate that members of the U.S. *artAB*-negative major clade do not have a significant advantage or disadvantage relative to closely related, Gifsy-1/*artAB*/*gogB*-harboring U.S. DT104 complex strains when exposed to stressors encountered during human and/or bovine digestion *in vitro*. However, ArtAB heterogeneity within the DT104 complex suggests clade-specific selection for or against maintenance of ArtAB. Thus, future studies querying the virulence characteristics of the U.S. *artAB*-negative major clade are needed.

*Data Summary:* Supplementary Data is available under DOI 10.5281/zenodo.7688792, with URL https://doi.org/10.5281/zenodo.7688792.

## INTRODUCTION

Prophages, which are viruses located within the genomes of bacteria, play important roles in the evolution of their microbial hosts [1–4]. In addition to possessing machinery that is antagonistic to host cell survival (e.g., virion production, lysis of host cells), many prophages encode accessory genes, which may provide the host with a selective advantage [1, 3, 5], including stress tolerance, resistance to antimicrobials and phage, biofilm formation, increased virulence, and evasion of the host immune system [1, 2, 4–8]. While they may persist within a lineage through vertical transmission [5, 6, 9], prophages can undergo gain and loss events within a population over time [1, 3]. Furthermore, integrated prophages can be hotspots for horizontal gene transfer (HGT) and genomic recombination, allowing their bacterial hosts to gain, lose, and exchange genetic information [4]. Thus, prophage-mediated HGT may confer novel functions, which allow the bacterial host to survive and compete in its environment, potentially contributing to the emergence of novel epidemic lineages [4, 10].

*Salmonella enterica* subsp. *enterica* serotype Typhimurium (*S.* Typhimurium) is among the *Salmonella* serotypes most commonly isolated from human and animal salmonellosis cases worldwide [11, 12] and is known to host a range of prophages within its chromosome [10]. Of particular concern is *S.* Typhimurium definitive type 104 (DT104), a lineage within *S.* Typhimurium that is known for its typical ampicillin-, chloramphenicol-, streptomycin-, sulfonamide-, and tetracycline-resistant (ACSSuT) phenotype, although its antimicrobial resistance (AMR) profile may vary [13]. Multidrug-resistant (MDR) DT104 is predicted to have emerged circa 1972 [13] and rapidly disseminated around the world in the following decades [13–15], culminating in a global epidemic among animals and humans in the 1990s [13–15]. However, despite its rapid global dissemination, DT104 does not appear to be more virulent than non-DT104 *S.* Typhimurium in a classical mouse model [16].

In addition to its characteristic MDR phenotype, DT104 is notable for its ability to produce ArtAB, a pertussis-like toxin that catalyzes ADP-ribosylation of host G proteins [17–19]. Treatment of various cell lines with purified ArtAB from DT104 recapitulates some of the phenotypes established for pertussis toxin cytotoxicity [20–22], such as the characteristic “cell clustering” phenotype in CHO-K1 cells [23], increased levels of intracellular cAMP in RAW 264.7 macrophage-like cells [18], and increased serum insulin levels (e.g., insulinemia); further, intraperitoneal injection of purified toxin in neonatal mice was fatal [18].

*artAB,* which encodes ArtAB, has been detected in representative strains of at least 88 *Salmonella* serotypes [24], and previous studies have found that *artAB* can be encoded by prophages (e.g., Gifsy-1, PhInv-1b) [17-19, 25-27]. Within *S.* Typhimurium specifically, *artAB* shares a strong association with DT104 relative to other *S.* Typhimurium lineages: while typically absent in most non-DT104 *S.* Typhimurium strains, the overwhelming majority of DT104 possess *artAB* [18]. In DT104 specifically, *artAB* has been identified within prophage Gifsy-1 (Figure 1A and Supplementary Figure S1) [10, 19, 25]. Gifsy-1 has been detected in numerous *Salmonella* serotypes [28] and has been shown to harbor virulence factors [29] such as *gogB*, *gipA*, and *gtgA* [29–31]; however, *artAB*-harboring Gifsy-1 has been proposed to be a characteristic feature of DT104 (Figure 1A and Supplementary Figure S1) [10].

**Figure 1.**
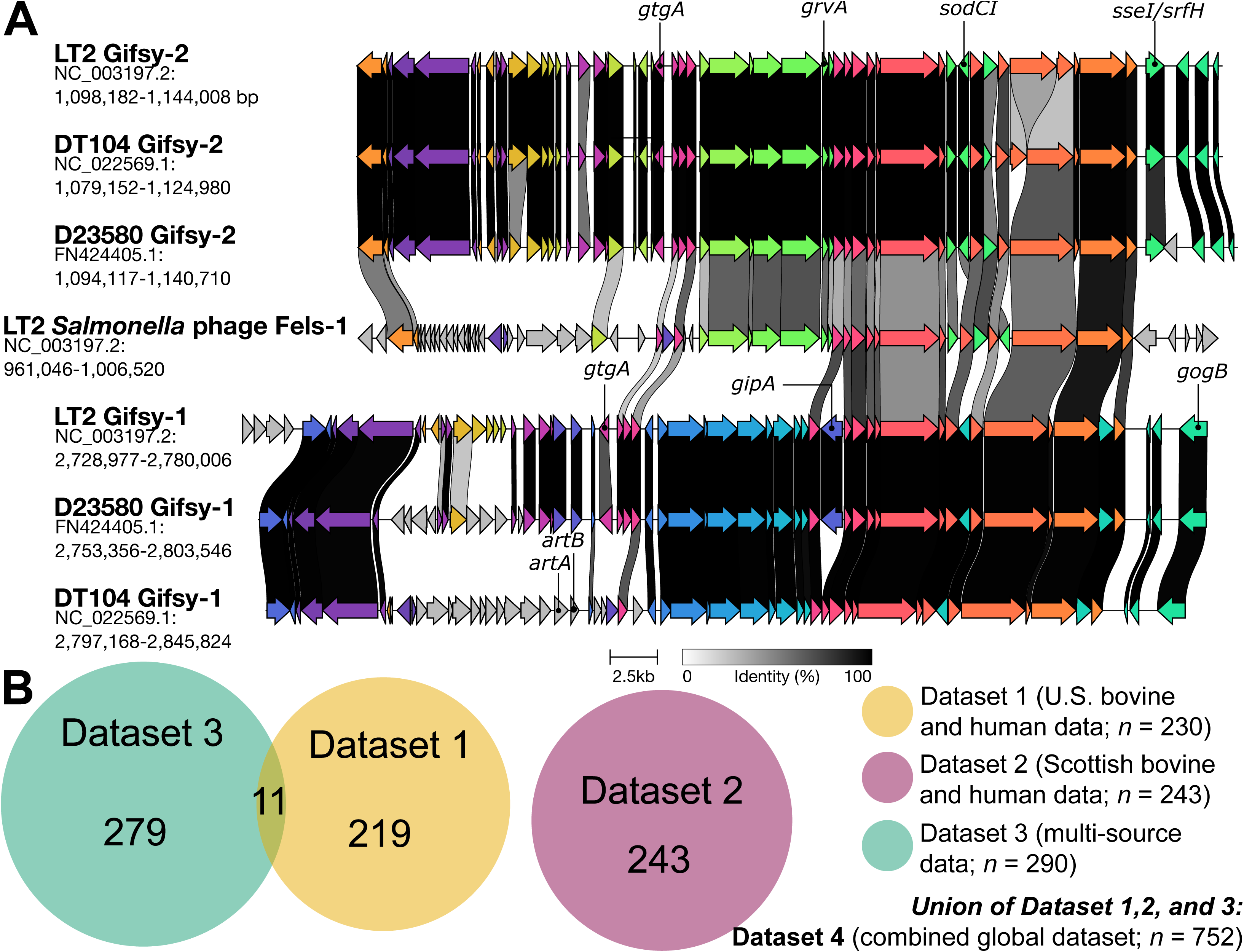
(A) Selected prophages that share homology with prophage Gifsy-1, as described in *Salmonella* Typhimurium strains (i) LT2, (ii) DT104, and (iii) D23580. Prophage regions were acquired from the PHASTER database and annotated using Prokka. clinker was used to compare prophage regions using default settings. Arrows correspond to open reading frames (ORFs), with grayscale links denoting the percent (%) amino acid identity shared between corresponding ORFs. Selected Gifsy-1 and Gifsy-2 virulence factors [29] are annotated. To view a similar plot constructed using all PHASTER prophage regions in LT2, DT104, and D23580, see Supplementary Figure S1. (B) Venn diagram showcasing the relationship between DT104 complex datasets used in this study. Numbers denote the number of genomes within a given dataset or subset of a dataset. For an extended version of this figure, see Supplementary Figure S2. For a flow chart with detailed descriptions of the datasets used in this study, see Supplementary Figure S3.

Despite their strong association, DT104 strains that lack *artAB* and, thus, the ability to produce ArtAB toxin, have been described on occasion (referred to hereafter as “*artAB*-negative” strains) [18, 32]. We identified three *artAB*-negative DT104 complex strains in a previous study of *S.* Typhimurium from cattle and humans in New York State (United States, U.S.) [32]. Because *artAB* tends to be prophage-encoded [17-19, 25, 26], we hypothesized that it may be possible for *artAB* to be gained or lost as an *artAB*-harboring prophage integrates or excises from a genome, or via HGT within an integrated prophage. However, the extent to which any of these scenarios occur is unknown. Using (i) 230 human- and bovine-associated DT104 complex genomes collected across the U.S., plus (ii) 752 DT104 complex genomes collected from a range of sources worldwide, we provide large-scale insight into the dynamics of *artAB* acquisition and loss within the DT104 complex.

## METHODS

### Acquisition of U.S. human- and bovine-associated DT104 complex genomic data and metadata

In a previous study of human- and bovine-associated *S.* Typhimurium from New York State, we identified three closely related, *artAB*-negative DT104 complex genomes from both humans and cattle (out of 14 total DT104 complex genomes from humans and cattle in New York State) [32]. Thus, as a first evaluation of *artAB* presence and absence in the DT104 complex, we compiled a set of DT104 complex genomes from humans and cattle across the U.S. (referred to hereafter as “Dataset 1 [U.S. bovine and human data]”; Figure 1B and Supplementary Figures S2 and S3).

To construct Dataset 1 (U.S. bovine and human data), we first collected genomic data derived from 14 human- and bovine-associated DT104 complex isolates from New York State, which we had sequenced in a previous study (members of the *S.* Typhimurium Lineage III cluster described in Supplementary Figures S2 and S5 of Carroll, et al.) [32]. We then aggregated these 14 New York State genomes with 223 human- and bovine-associated DT104 complex genomes from across the U.S., as described previously [32]. Briefly, paired-end Illumina short reads associated with 223 *S.* Typhimurium genomes meeting the following criteria were downloaded from the National Center for Biotechnology Information (NCBI) Sequence Read Archive (SRA; https://www.ncbi.nlm.nih.gov/sra, accessed November 29, 2018), using accession numbers provided by Enterobase and the SRA Toolkit version 2.9.3 [33–36]: (i) genomes were serotyped as *S.* Typhimurium *in silico* using the implementation of SISTR [37] in Enterobase; (ii) the country of isolation was the United States; (iii) the isolation source was reported as either “Human” or “Bovine” in the “Source Niche” and “Source Type” fields in Enterobase, respectively; (iv) genomes had an isolation year reported in Enterobase; (v) using RhierBAPS [38], genomes were assigned to the DT104 complex, a well-supported cluster within the larger bovine- and human-associated U.S. *S.* Typhimurium phylogeny, which clustered among known DT104 genomes from other countries (see Supplementary Figures S2 and S5 of Carroll, et al.) [32].

Trimmomatic version 0.36 [39] was used to trim low quality bases and Illumina adapters from all read sets using the default settings for paired-end reads, and SPAdes version 3.13.0 [40] was used to assemble all genomes using default settings plus the “careful” option. FastQC version 0.11.5 [41] and QUAST version 4.0 [42] were used to assess the quality of each read pair set and assembly, respectively, and MultiQC version 1.6 [43] was used to aggregate all FastQC and QUAST results. Trimmed paired-end read sets/assemblies that were flagged by MultiQC as meeting any of the following conditions were excluded: (i) Illumina adapters present after trimming (*n* = 2), (ii) an abnormal per sequence GC content distribution (*n* = 3), (iii) an assembly with over 200 contigs (*n* = 11), and (iv) a sequence quality histogram flagged as poor quality (*n* = 2). After excluding genomes that met these conditions, a set of 219 DT104 complex genomes was produced (Figure 1B, Supplementary Figures S2 and S3).

Finally, the 219 U.S. human- and bovine-associated DT104 complex genomes identified here were supplemented with 11 U.S. bovine- and human-associated DT104 genomes from a previous study [13], which did not have metadata available in Enterobase at the time and were thus not included in the initial set of 219 bovine- and human-associated U.S. DT104 complex genomes. Overall, the search conducted here produced a set of 230 bovine- and human-associated U.S. DT104 complex genomes, which were used in subsequent steps (i.e., Dataset 1 [U.S. bovine and human data]; Figures 1B and 2A, Supplementary Figures S2 and S3, and Supplementary Tables S1 and S2).

**Figure 2.**
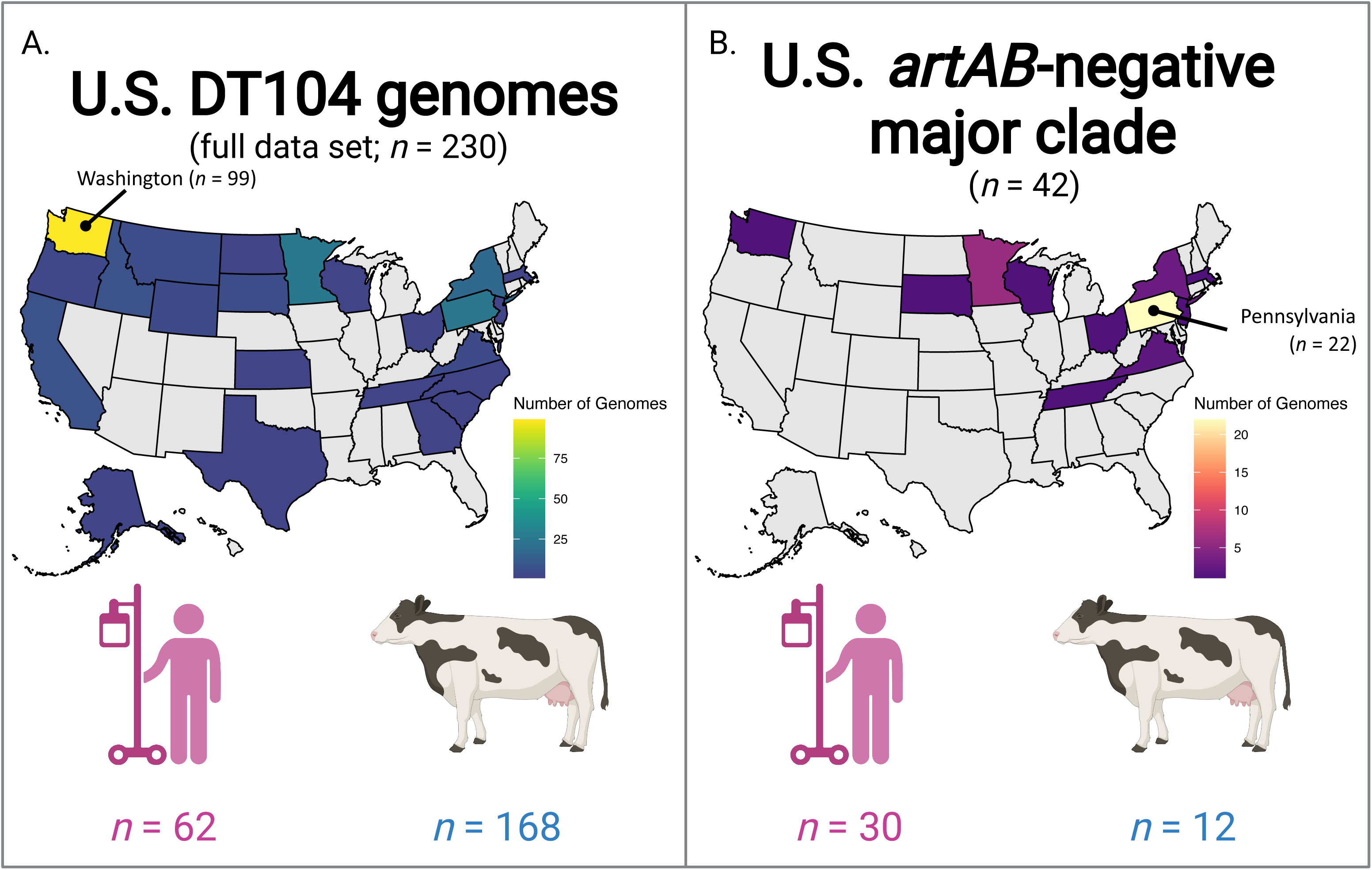
Geographic and source origins (i.e., human or bovine) of (A) all 230 human- and bovine-associated United States (U.S.) DT104 complex genomes queried in this study (i.e., Dataset 1 [U.S. bovine and human data]), and (B) 42 Gifsy-1/*artAB*/*gogB*-negative genomes assigned to the U.S. *artAB*-negative major clade. U.S. states shown in gray did not contribute any genomes to the respective data set. The U.S. state that contributed the most genomes to its respective data set is labeled. The figure was created using BioRender (https://biorender.com/) and the “plot_usmap” function in the usmap version 0.6.0 R package [146].

### *In silico* detection of prophage, antimicrobial resistance genes, plasmid replicons, and virulence factors

To identify putative prophage regions in all 230 genomes in Dataset 1 (U.S. bovine and human data), each assembly was submitted to the PHASTER web server (https://phaster.ca/) via the URL API [44, 45], with the “contigs” option set to “1” (Supplementary Table S3). To compare prophage regions identified in Dataset 1 (U.S. bovine and human data) genomes to previously described prophage in well-characterized *S.* Typhimurium strains, prophage in the following *S.* Typhimurium strains were obtained from the PHASTER prophage database (accessed September 18, 2020): (i) LT2 (NCBI Nucleotide accession NC_003197.2), (ii) DT104 (NCBI Nucleotide accession NC_022569.1), (iii) D23580 (NCBI Nucleotide accession FN424405.1), and (iv) SL1344 (NCBI Nucleotide accession NC_016810.1; Figure 1A and Supplementary Figure S1). All prophage regions were annotated using Prokka version 1.14.6 [46], using default settings and the “Viruses” kingdom database. The resulting GFF and FNA files produced by Prokka were supplied to clinker version 0.0.26 [47], which was used to perform pairwise alignments of all genes within prophage regions using default settings.

ABRicate version 0.8 [48] was used to detect antimicrobial resistance (AMR) genes, plasmid replicons, and virulence factors in each assembled DT104 complex genome using NCBI’s National Database of Antibiotic Resistant Organisms (NDARO) [49], the PlasmidFinder database [50], and the Virulence Factor Database (VFDB) [51], respectively, using minimum nucleotide identity and coverage thresholds of 75 and 50%, respectively (all databases accessed December 10, 2020; Supplementary Table S3). The aforementioned ABRicate analyses were repeated, using a minimum coverage threshold of 0% (e.g., to confirm that virulence factors discussed in the manuscript were absent from genomes in which they were not initially detected).

Each assembled genome was additionally queried for the presence of selected virulence factors, which have previously been associated with prophage in *Salmonella* [29]: (i) *artAB* (NCBI Nucleotide accession AB104436.1), (ii) *gogA* (European Nucleotide Archive [ENA] accession EAA7850902.1), (iii) *gtgA* (ENA accession PVI70081.1), and (iv) *gipA* (ENA accession CAI93790.1). Assembled genomes were queried for selected virulence factors using the command-line implementation of nucleotide BLAST (blastn) version 2.11.0 [52], using default settings plus a minimum coverage threshold of 40% (Supplementary Table S3). To confirm that the aforementioned genes were absent from genomes in which they were not initially detected, all genomes were queried again (i) as described above, with the coverage threshold lowered to 0%; and (ii) using translated nucleotide BLAST (tblastx; Supplementary Tables S4 and S5). ARIBA version 2.14.6 [53] was used to further confirm *artAB* and *gogB* presence/absence in all genomes with associated paired-end Illumina reads (Supplementary Table S6 and Supplementary Text).

### Variant calling and maximum likelihood phylogeny construction within Dataset 1 (U.S. bovine and human data)

Core single nucleotide polymorphisms (SNPs) were identified among all 230 genomes within Dataset 1 (U.S. bovine and human data), using the default pipeline implemented in Snippy version 4.6.0 (https://github.com/tseemann/snippy; Supplementary Tables S1 and S2 and Supplementary Text) [54–67]. The closed DT104 chromosome (NCBI Nucleotide accession NC_022569.1) was used as a reference, and core SNPs identified in regions of the DT104 chromosome predicted to belong to phage were masked (Supplementary Text). Gubbins version 2.4.1 [68] was used to identify and remove recombination events in all genomes using default settings, and snp-sites was used to query the resulting recombination-free alignment for core SNPs (i.e., using the “-c” option).

A maximum likelihood (ML) phylogeny was constructed with IQ-TREE version 1.5.4 [69], using (i) the resulting core SNPs as input, (ii) the optimal nucleotide substitution model selected using ModelFinder [70, 71], (iii) an ascertainment bias correction to account for the use of solely variant sites, and (iv) 1,000 replicates of the ultrafast bootstrap approximation (Supplementary Text) [72, 73]. TempEst version 1.5.3 [74] was used to assess the temporal structure of the resulting unrooted ML phylogeny, using the best-fitting root and the *R*^2^ function (*R*^2^ = 0.33, slope = 3.05×10^-7^ substitutions/site/year, X-intercept = 1988.1). The unrooted ML phylogeny was additionally rooted and time scaled using LSD2 version 1.4.2.2 [75], using tip dates corresponding to the year of isolation reported for each genome (Supplementary Text). The resulting rooted, time-scaled ML phylogeny was viewed using FigTree version 1.4.4 [76] (Supplementary Data).

### Dataset 1 (U.S. bovine and human data) Bayesian time-scaled phylogeny construction

In addition to constructing a time-scaled ML phylogeny (see section “Variant calling and maximum likelihood phylogeny construction within Dataset 1 [U.S. bovine and human data]” above), a Bayesian approach was additionally employed to construct a time-scaled phylogeny, using a subset of 146 Dataset 1 (U.S. bovine and human data) genomes (Supplementary Figure S4 and Supplementary Text). All aforementioned SNP calling and ML phylogeny construction steps were repeated within the 146-genome Dataset 1 (U.S. bovine and human data) subset, and the resulting ML phylogeny was time-scaled using TempEst and LSD2 as described above (see section “Variant calling and maximum likelihood phylogeny construction within Dataset 1 [U.S. bovine and human data]” above; Supplementary Table S7, Supplementary Data, and Supplementary Text).

BEAST2 version 2.5.1 [77, 78] was used to construct a tip-dated phylogeny, using core SNPs detected among the 146-genome Dataset 1 (U.S. bovine and human data) subset as input (Supplementary Text). An initial clock rate of 2.79×10^-7^ substitutions/site/year [13] was used, along with an ascertainment bias correction to account for the use of solely variant sites [79]. bmodeltest [80] was used to infer a substitution model using Bayesian model averaging, with transitions and transversions split. A relaxed lognormal molecular clock [81] and a coalescent Bayesian skyline population model [82] were used, as these models have been selected as the optimal clock/population model combination for DT104 previously [13] (Supplementary Text). Five independent BEAST2 runs (i.e., BEAST2 runs with different random seeds) were performed, using chain lengths of at least 100 million generations, sampling every 10 thousand generations. LogCombiner-2 was used to aggregate the resulting log and tree files with 10% of the states treated as burn-in, and TreeAnnotator-2 was used to produce a maximum clade credibility (MCC) tree using Common Ancestor node heights (Supplementary Figure S5, Supplementary Table S8, and Supplementary Data). The resulting phylogenies were displayed and annotated using R version 4.1.2 (Figures 3 and 4, Supplementary Figures S6-S9, and Supplementary Text) [83–88].

**Figure 3.**
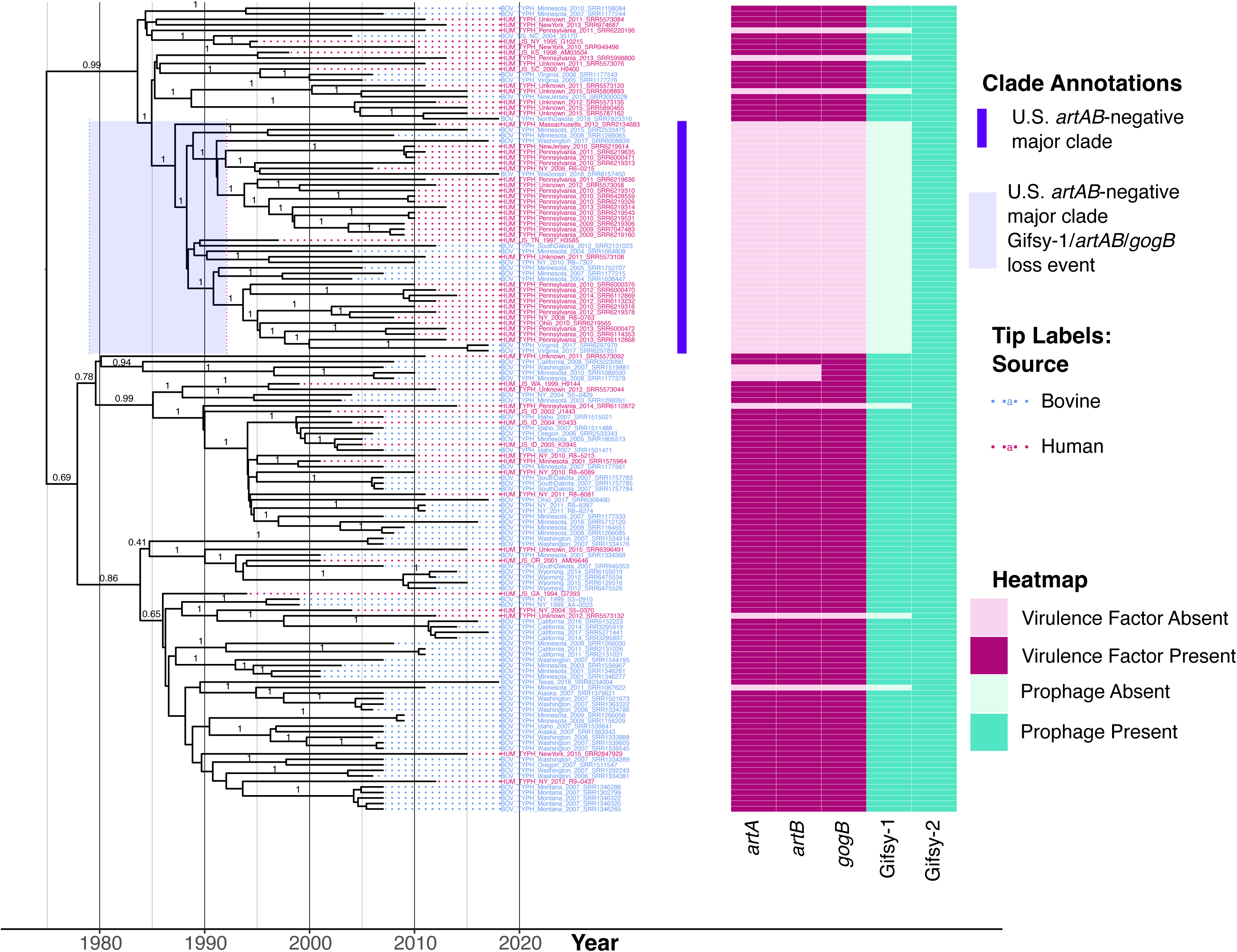
Bayesian time-scaled phylogeny constructed using 146 human- and bovine-associated DT104 complex genomes collected in the United States (U.S.; i.e., a subset of genomes from Dataset 1 [U.S. bovine and human data]). Tip label colors denote the isolation source reported for each genome (human or bovine in pink and blue, respectively). The heatmap to the right of the phylogeny denotes the presence and absence of (i) selected virulence factors (dark and light pink, respectively) and (ii) prophage (dark and light green, respectively). The U.S. *artAB*-negative major clade is denoted by the bright purple bar; light purple shading around the node of the U.S. *artAB*-negative major clade denotes the 95% highest posterior density (HPD) interval, in which Gifsy-1/*artAB*/*gogB* were predicted to have been lost. The phylogeny was constructed and rooted using BEAST2. Time in years is plotted along the X-axis, while branch labels correspond to posterior probabilities of branch support (selected for readability). For extended versions of this figure, see Supplementary Figures S6-S8.

**Figure 4.**
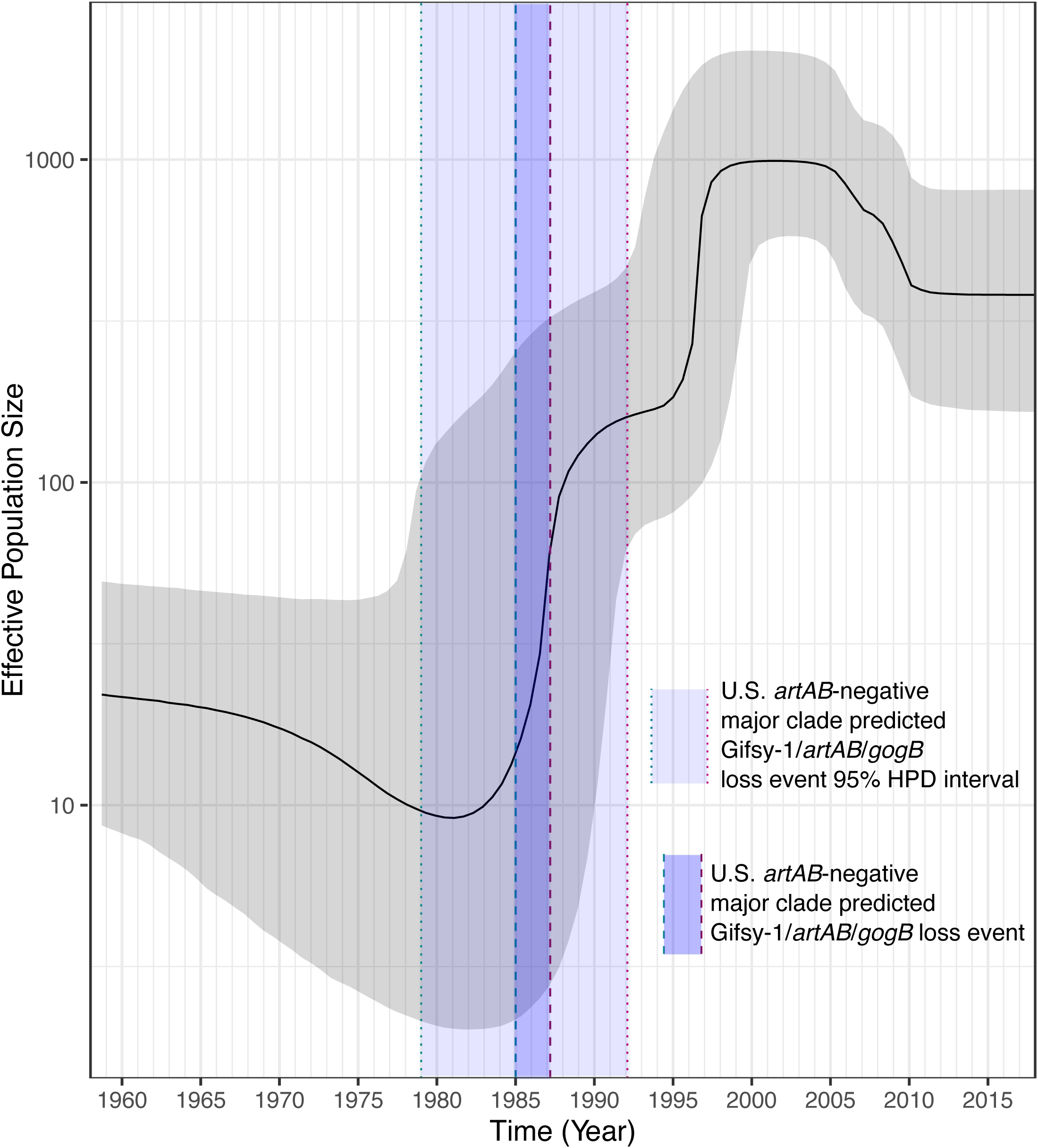
Coalescent Bayesian Skyline plot constructed using 146 U.S. bovine- and human-associated *S.* Typhimurium DT104 complex genomes (i.e., a subset of genomes from Dataset 1 [U.S. bovine and human data]). Effective population size and time in years are plotted along the Y-and X-axes, respectively. The median effective population size estimate is denoted by the solid black line, with upper and lower 95% highest posterior density (HPD) interval bounds denoted by gray shading. The interval shaded in light blue and bounded by dashed vertical lines denotes the time interval in which Gifsy-1/*artAB*/*gogB* were predicted to have been lost by the common ancestor of the U.S. *artAB*-negative major clade (corresponding to the years 1985.0 and 1987.2, denoted by turquoise and pink dashed lines, respectively). The dotted turquoise and pink vertical lines correspond to the 95% HPD interval lower and upper bounds for Gifsy-1/*artAB*/*gogB* loss among members of the U.S. *artAB*-negative major clade (corresponding to the years 1979.0 and 1992.1, respectively).

### *artAB* ancestral state reconstruction for Dataset 1 (U.S. bovine and human data)

To estimate ancestral character states of internal nodes in the Dataset 1 (U.S. bovine and human data) phylogeny as they related to *artAB* presence/absence (i.e., whether a node in the tree represented an ancestor that was more likely to be *artAB*-positive or *artAB*-negative), the presence or absence of *artAB* within each genome was treated as a binary state (see section “*In silico* detection of prophage, antimicrobial resistance genes, plasmid replicons, and virulence factors” above). *artAB* ancestral state reconstruction runs were performed using the BEAST2 time-scaled Bayesian Dataset 1 (U.S. bovine and human data) phylogeny as input (*n* = 146 genomes; see section “Dataset 1 [U.S. bovine and human data] Bayesian time-scaled phylogeny construction” above; Supplementary Data and Supplementary Text). Stochastic character maps were simulated on the phylogeny using the make.simmap function in the phytools version 1.0-1 R package [89] and the all-rates-different (ARD) model in the ape version 5.6-1 package [90, 91]; two root node priors were tested (Supplementary Text). The resulting phylogenies (one for each root node prior) were plotted using the densityMap function in the phytools R package (Supplementary Figures S10-S12, Supplementary Data).

### Pan-genome characterization of Dataset 1 (U.S. bovine and human data)

Prokka version 1.13.3 [46] was used to annotate all 230 genomes within Dataset 1 (U.S. bovine and human data), using the “Bacteria” database and default settings (Supplementary **Tables S1** and S2). GFF files produced by Prokka were supplied as input to Panaroo version 1.2.7 [92], which was used to identify core- and pan-genome orthologous gene clusters among the 230 Dataset 1 (U.S. bovine and human data) genomes (Supplementary Text) [93, 94]. The LSD2 time-scaled ML phylogeny for Dataset 1 (U.S. bovine and human data) (see section “Variant calling and maximum likelihood phylogeny construction within Dataset 1 [U.S. bovine and human data]” above) was supplied as input to Panaroo’s “panaroo-img” and “panaroo-fmg” commands, which were used to estimate the pan-genome size under the Infinitely Many Genes (IMG) [95, 96] and Finite Many Genes (FMG) models (with 100 bootstrap replicates) [97], respectively (Supplementary **Figure S13**).

Reference pan-genome coding sequences (CDS) identified by Panaroo underwent functional annotation using the eggNOG-mapper version 2 webserver (http://eggnog-mapper.embl.de/; accessed July 24, 2022) using default settings [98, 99]. The “table” function in R was used to identify genes associated with (i) prophage Gifsy-1 presence/absence (Supplementary Table S9) and (ii) clade membership (Supplementary Table S10); the “fisher.test” function in R’s stats package was used to conduct two-sided Fisher’s exact tests, and the “p.adjust” function was used to control the false discovery rate (FDR; i.e., p.adjust method = “fdr”) [100].

### Genome-wide identification of host-associated orthologous gene clusters for Dataset 1 (U.S. bovine and human data)

The treeWAS version 1.0 R package [101] was used to identify potential orthologous gene cluster-host associations among the 230 human- and bovine-associated U.S. DT104 complex genomes in Dataset 1 (U.S. bovine and human data) (i.e., whether an orthologous gene cluster identified with Panaroo was human- or bovine-associated while accounting for population structure; Supplementary Text). No orthologous gene clusters were found to be significantly associated with isolation source via any of the treeWAS association tests (FDR-corrected *P*-value > 0.10).

### Acquisition of global DT104 complex genomic data and metadata

To compare the 230 U.S. human- and bovine-associated DT104 complex genomes in Dataset 1 (U.S. bovine and human data) to a larger set of DT104 complex genomes from numerous sources worldwide, genomic data associated with the following studies were downloaded via Enterobase: (i) 243 bovine- and human-associated DT104 isolates from a study of between-host transmission within Scotland [102] (referred to hereafter as “Dataset 2 [Scottish bovine and human data]”; Supplementary Text); (ii) 290 DT104 isolates from a variety of sources and countries from a study describing the global spread of DT104 [13] (referred to hereafter as “Dataset 3 [multi-source data]”; eleven of the 290 genomes were isolated from cattle and humans in the U.S. and thus had also been included in Dataset 1 [U.S. bovine and human data], Figure 1B, Supplementary Figures S2 and S3, Supplementary Table S1, and Supplementary Text).

The following datasets were aggregated to create a final set of 752 DT104 complex genomes derived from numerous countries and isolation sources, which was used in subsequent steps (referred to hereafter as “Dataset 4 [combined global dataset]”; Figure 1B, Supplementary Figures S2 and S3, and Supplementary Tables S1 and S2): (i) Dataset 1 (U.S. bovine and human data) (*n* = 230 DT104 complex genomes), (ii) Dataset 2 (Scottish bovine and human data) (*n* = 243 DT104 genomes), and (iii) Dataset 3 (multi-source data) (*n* = 290 DT104 genomes, including 11 genomes that were part of Dataset 1 [U.S. bovine and human data]). QUAST version 4.5 was used to assess the quality of all 752 genomes in Dataset 4 (combined global dataset) (Supplementary Tables S1 and S2). Prophage, AMR genes, plasmid replicons, and virulence factors were detected in all 752 Dataset 4 (combined global dataset) genomes as described above (see section “*In silico* detection of prophage, antimicrobial resistance genes, plasmid replicons, and virulence factors” above; Supplementary Tables S3-S5).

### Variant calling and maximum likelihood phylogeny construction within Dataset 4 (combined global dataset)

To identify core SNPs present in all 752 DT104 complex genomes within Dataset 4 (combined global dataset), Parsnp and HarvestTools version 1.2 [103] were used, as Parsnp easily scales to large data sets (Supplementary Tables S1 and S2) [103]. Assembled genomes were used as input for Parsnp, along with the closed DT104 chromosome as a reference (NCBI Nucleotide accession NC_022569.1) and Parsnp’s implementation of PhiPack [104] to filter recombination.

Core SNPs detected among all 752 assembled genomes within Dataset 4 (combined global dataset) were supplied as input to IQ-TREE version 1.5.4, which was used to construct a ML phylogeny as described above; the resulting ML phylogeny was rooted and time-scaled using LSD2 as described above (see section “Variant calling and maximum likelihood phylogeny construction within Dataset 1 [U.S. bovine and human data]” above; Supplementary Data and Supplementary Text). The resulting LSD2 time-scaled ML phylogeny was annotated using the Interactive Tree of Life (iTOL) version 6 webserver (https://itol.embl.de/, accessed March 7, 2022; Figure 5, Supplementary Figure S14, and Supplementary Data) [105]. The LSD2 time-scaled ML phylogeny for Dataset 4 (combined global dataset) was further used for *artAB* presence/absence ancestral state reconstruction as described above (see section “*artAB* ancestral state reconstruction for Dataset 1 [U.S. bovine and human data]” above; Supplementary Figure S15).

**Figure 5.**
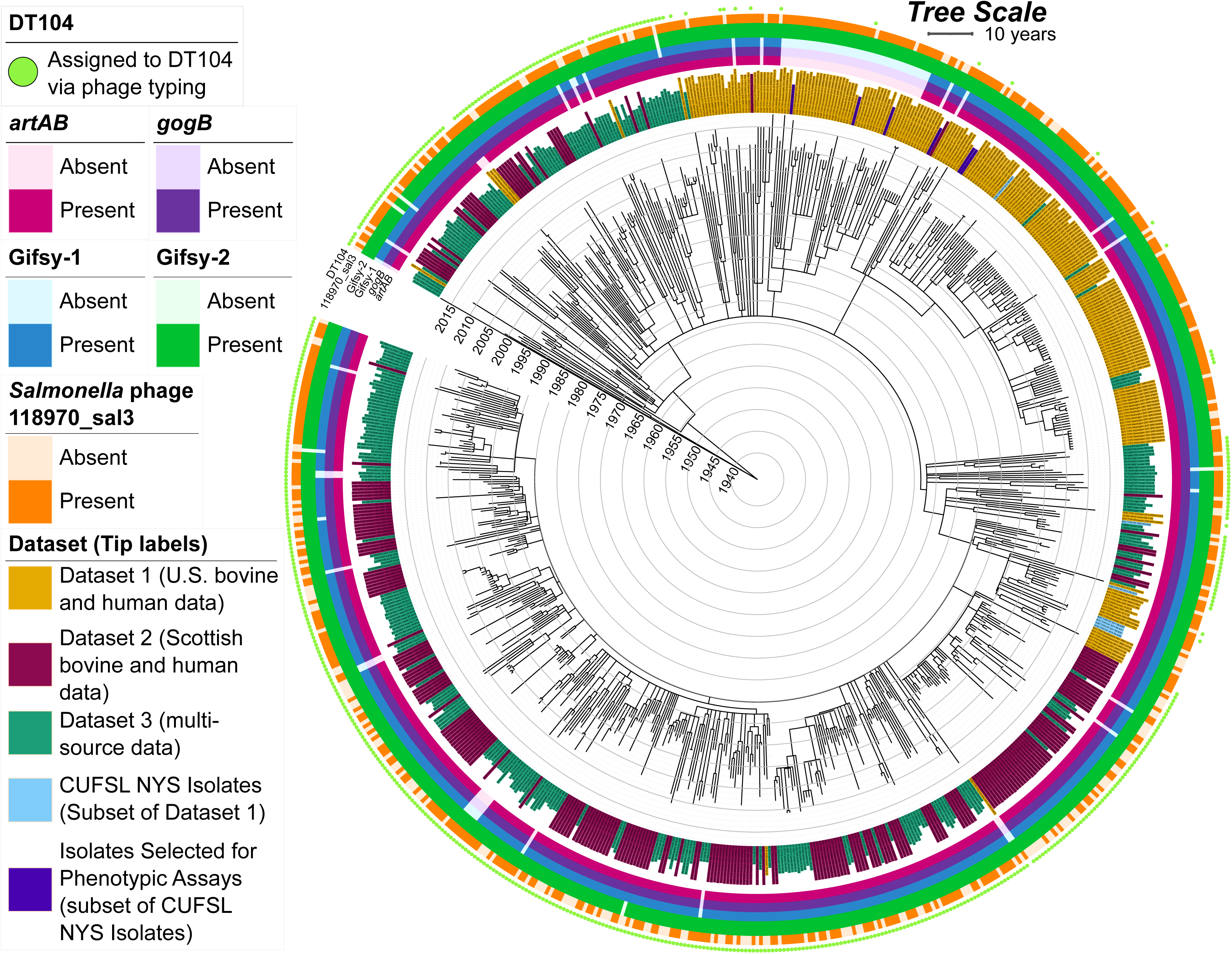
Time-scaled maximum likelihood (ML) phylogeny constructed using 752 DT104 complex genomes (i.e., Dataset 4 [combined global dataset], the union of Dataset 1 [U.S. bovine and human data], Dataset 2 [Scottish bovine and human data], and Dataset 3 [multi-source data]). Tip label colors denote the dataset or dataset subset with which each genome is affiliated (“Dataset”; eleven genomes within the intersection of Dataset 1 [U.S. bovine and human data] and Dataset 3 [multi-source data] were colored as Dataset 1 [U.S. bovine and human data]). The heatmap encompassing the phylogeny denotes the presence and absence of selected virulence factors and intact prophage (identified via nucleotide BLAST [blastn; default settings, with no minimum identity or coverage threshold employed] and PHASTER, respectively). Bright green circles in the outermost ring surrounding the phylogeny denote genomes reportedly assigned to DT104 using phage typing (“DT104”). The ML phylogeny was constructed using IQ-TREE and rooted and time-scaled using LSD2. Branch lengths are reported in years. For an extended version of this figure, see Supplementary Figure S14.

### Pan-genome characterization of Dataset 4 (combined global dataset)

Pan-genome analyses were carried out for Dataset 4 (combined global dataset) as described above (see section “Pan-genome characterization of Dataset 1 [U.S. bovine and human data]” above; Supplementary Tables S1 and S2 and Supplementary Text). The pan-genome size for Dataset 4 (combined global dataset) was estimated using Panaroo’s “panaroo-img” and “panaroo-fmg” commands, using the LSD2 time-scaled ML phylogeny for Dataset 4 (combined global dataset) as input (see section “Variant calling and maximum likelihood phylogeny construction within Dataset 4 [combined global dataset]” above; Supplementary Figure S13).

### Strain selection for phenotypic stress assays

Phenotypic stress assays (discussed in detail in the sections below) were used to compare (i) bovine- and human-associated, Gifsy-1/*artAB*/*gogB*-positive U.S. DT104 complex strains to (ii) bovine- and human-associated, Gifsy-1/*artAB*/*gogB*-negative U.S. DT104 complex strains. Thus, additional analyses were performed to identify the most closely related, Gifsy-1/*artAB*/*gogB*-positive and -negative strains available in the Cornell University Food Safety Laboratory (CUFSL) culture collection for phenotypic testing (Supplementary Figures S2 and S3, Supplementary Table S2, and Supplementary Text) [106–108]. Overall, the CUFSL DT104 complex genomes differed little in terms of their core and pan-genome compositions (Supplementary Figure S16 and Supplementary Text).

Considering both (i) core- and pan-genome similarities between all 13 available CUFSL DT104 complex genomes (Supplementary Figure S16 and Supplementary Table S2), as well as (ii) Gifsy-1/*artAB*/*gogB* presence and absence, we selected six closely related, CUFSL DT104 complex strains to undergo phenotypic characterization: three Gifsy-1/*artAB*/*gogB*-positive strains, and three Gifsy-1/*artAB*/*gogB*-negative strains (Figure 5, Supplementary Figure S16, Supplementary Table S11, and Supplementary Text). All three Gifsy-1/*artAB*/*gogB*-negative strains were members of the U.S. *artAB*-negative major clade (discussed in detail in the “Results” section below; Figure 5 and Supplementary Table S11). All six selected strains had been isolated from humans or cattle in New York State (U.S.) and were part of Dataset 1 (U.S. bovine and human data) (Supplementary Figures S2 and S3 and Supplementary Tables S2 and S11).

### Phenotypic assays

Six carefully selected, human- and bovine-associated DT104 complex strains from New York State (U.S.), which were available to us in the CUFSL culture collection, were characterized using phenotypic assays (see section “Strain selection for phenotypic stress assays” above; Supplementary Table S11). All strain stocks were maintained in CRYOBANK^®^ tubes (Mast Ltd., Reinfeld, Germany) at -80°C. Strains were streaked out from stocks on tryptic soy agar (TSA; Merck KGaA, Darmstadt, Germany) and incubated overnight at 37°C. Single colonies from those plates were inoculated in 5 mL of tryptic soy broth (TSB; Merck KGaA, Darmstadt, Germany) and incubated for 16 - 18 h at 37°C with shaking at 200 rpm. The resulting overnight cultures were diluted 1/100 into 5 mL of fresh, pre-warmed TSB, followed by incubation at 37°C with shaking at 200 rpm to allow cultures to reach mid log phase (defined as OD_600_ of 0.4; 1-2 x10^8^ CFU/mL). These cultures were used as input into three different phenotypic assays (exposure to ruminal fluid, acid stress, and bile stress; discussed in detail below). Bacterial enumeration before and after stress exposure was performed by direct colony counts of tilt plates according to Kühbacher et al. [109].

To evaluate exposure to ruminal fluid (RF), approximately 2 L of RF was acquired from a Jersey cow with a ruminal fistula on each experimental day prior to the experiments (same collection time was used for each experiment). The RF was immediately filtered through a cellulose filter (Labsolute® Type 80, Th. Geyer GmbH& Co. KG., Renningen, Germany) to remove any large debris, and the pH was measured, ranging from 7.20 to 7.62. Mid-log phase cultures were prepared and inoculated into the RF at two different concentrations. One hundred µl of culture suspensions were inoculated into 5mL of the RF at final concentrations of 10^8^ (high) and 10^5^ (low) CFU/mL and incubated for 1 h at 37°C without shaking with enumeration by direct colony counting on XLT-4 agar (Oxoid Ltd., Basingstoke, UK) prior and after RF exposure (Supplementary Table S12). The absence of *Salmonella* in the RF at the start of the experiments was confirmed by plating on XLT-4 agar.

Acid stress resistance of the different strains at pH 3.5 with and without prior adaption was tested using an adopted protocol from Horlbog et al. [110]. To carry out the acid stress assay, the pH of the TSB was adjusted with hydrochloric acid solution (1M and 6 M HCL; Merck KGaA, Darmstadt, Germany) immediately prior to the experiment. 1 mL aliquots of mid log phase cultures were transferred to reaction tubes and centrifuged at 14,000 x g for 10 min. For the non-adapted acid stress experiments, the pellets were resuspended in 1mL TSB pH 3.5 and incubated for 1 h at 37°C without shaking. For acid adaption, 1 mL of the same cultures were pelleted, resuspended in 1 mL TSB adjusted to pH 5.5 and incubated for 1h at 37°C (without shaking). Afterwards, the cultures were centrifuged again, resuspended in 1mL TSB pH 3.5, and incubated for 1 h at 37°C without shaking. Bacteria enumeration was performed before and after the one-hour incubation at pH 3.5 (Supplementary Table S13).

Susceptibility to bile salts (cholic acid and deoxycholic acid in a mixture of 1:1, Bile Salts No.3, Thermo Fisher Scientific Inc., Waltham, USA) was tested in two different concentrations: 14.5 mmol/L corresponding to 0.6% [111] and 26.0 mmol/L corresponding to 1.1% [112] were chosen to represent reasonable physiological states in the duodenum. Bile salts were added, and the pH of the TSB was adjusted to 5.5 (TSB-bile) immediately prior to the experiment. Mid log phase cultures were centrifuged, resuspended in TSB-bile, incubated for 1h 37°C without shaking, and enumerated by direct colony counting prior and after bile exposure (Supplementary Table S14).

For each stress assay, base-ten logarithmic fold change (FC) values were calculated as follows: FC = log CFU/g at the start of the experiments – log CFU/g after the stress assay. Analysis of Variance (ANOVA) for the interpretation of the phenotypic assays were conducted using the “aov” function in R’s “stats” package, with the FC values for the respective assay treated as a response. Figures were designed using the ggplot2 package.

### Data availability

Strain metadata, genome quality metrics, and Enterobase accession numbers for all publicly available genomes queried in this study are available in Supplementary Table S1. Strain metadata, genome quality metrics, CUFSL IDs [106], and NCBI BioSample accession numbers [113] for the 13 New York State CUFSL DT104 complex strains queried in this study (including those queried via phenotypic assays) are available in Supplementary Table S2. LSD2 results (for Dataset 1 [U.S. bovine and human data] and Dataset 4 [combined global dataset]) and BEAST2 results (for subsets of Dataset 1 [U.S. bovine and human data]) are available as Supplementary Data.

## RESULTS

### Human- and bovine-associated DT104 complex strains from the U.S. harbor *artAB* on prophage Gifsy-1

Within the set of 230 human- and bovine-associated U.S. DT104 complex genomes (i.e., Dataset 1 [U.S. bovine and human data]; Figure 2A) [32], *artAB* was present in over 75% of genomes (177 of 230, 77.0%; Figure 3, Table 1, and Supplementary Figures S6-S8). *artAB* presence and absence was strongly associated with the presence and absence of anti-inflammatory effector *gogB* (two-sided Fisher’s Exact Test [FET] raw *P*-value < 2.2×10^-16^, infinite odds ratio [OR]), as co-occurrence was observed in all 177 *artAB*-harboring Dataset 1 (U.S. bovine and human data) genomes (100.0%; Figure 3, Table 1, and Supplementary Figures S6-S8). Additionally, within Dataset 1 (U.S. bovine and human data), *artAB* and *gogB* presence was strongly associated with the presence of prophage Gifsy-1 (NCBI Nucleotide accession NC_010392.1; two-sided FET raw *P*-value < 2.2×10^-16^, infinite OR; Figure 3, Table 1, and Supplementary Figures S6-S8).

**Table 1.**
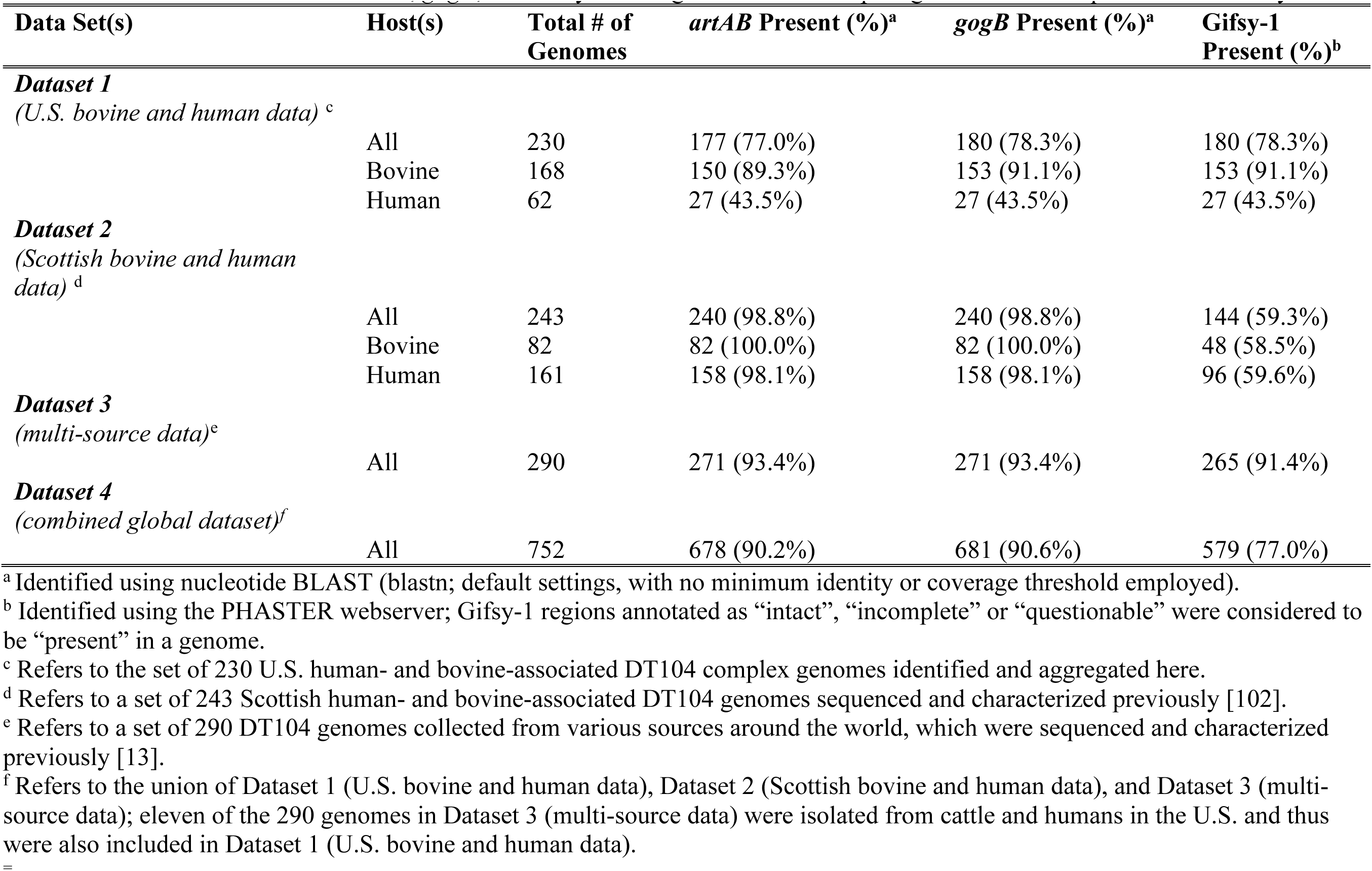
Presence and absence of artAB, gogB, and Gifsy-1 among four DT104 complex genome datasets queried in this study.

| Data Set(s) | Host(s) | Total # of Genomes | *artAB* Present (%)<sup>a</sup> | *gogB* Present (%)<sup>a</sup> | Gifsy-1 Present (%)<sup>b</sup> |
| --- | --- | --- | --- | --- | --- |
| **Dataset 1**<br>(U.S. bovine and human data)<sup>c</sup> |  |  |  |  |  |
|  | All | 230 | 177 (77.0%) | 180 (78.3%) | 180 (78.3%) |
|  | Bovine | 168 | 150 (89.3%) | 153 (91.1%) | 153 (91.1%) |
|  | Human | 62 | 27 (43.5%) | 27 (43.5%) | 27 (43.5%) |
| **Dataset 2**<br>(Scottish bovine and human data)<sup>d</sup> |  |  |  |  |  |
|  | All | 243 | 240 (98.8%) | 240 (98.8%) | 144 (59.3%) |
|  | Bovine | 82 | 82 (100.0%) | 82 (100.0%) | 48 (58.5%) |
|  | Human | 161 | 158 (98.1%) | 158 (98.1%) | 96 (59.6%) |
| **Dataset 3**<br>(multi-source data)<sup>e</sup> |  |  |  |  |  |
|  | All | 290 | 271 (93.4%) | 271 (93.4%) | 265 (91.4%) |
| **Dataset 4**<br>(combined global dataset)<sup>f</sup> |  |  |  |  |  |
|  | All | 752 | 678 (90.2%) | 681 (90.6%) | 579 (77.0%) |

<sup>a</sup> Identified using nucleotide BLAST (blastn; default settings, with no minimum identity or coverage threshold employed).
<sup>b</sup> Identified using the PHASTER webserver; Gifsy-1 regions annotated as “intact”, “incomplete” or “questionable” were considered to be “present” in a genome.
<sup>c</sup> Refers to the set of 230 U.S. human- and bovine-associated DT104 complex genomes identified and aggregated here.
<sup>d</sup> Refers to a set of 243 Scottish human- and bovine-associated DT104 genomes sequenced and characterized previously [102].
<sup>e</sup> Refers to a set of 290 DT104 genomes collected from various sources around the world, which were sequenced and characterized previously [13].
<sup>f</sup> Refers to the union of Dataset 1 (U.S. bovine and human data), Dataset 2 (Scottish bovine and human data), and Dataset 3 (multi-source data); eleven of the 290 genomes in Dataset 3 (multi-source data) were isolated from cattle and humans in the U.S. and thus were also included in Dataset 1 (U.S. bovine and human data).

Subsequent investigation confirmed that, for all 177 *artAB-*harboring Dataset 1 (U.S. bovine and human data) genomes, *artAB* was located within Gifsy-1 prophage regions classified as “intact” via PHASTER (Table 2, Supplementary Figures S17 and S18, and Supplementary Table S5). *gogB* was largely harbored within regions annotated via PHASTER as Gifsy-1 (126 of 180 *gogB*-harboring Dataset 1 [U.S. bovine and human data] genomes, 70.0%), although only 51 of these Gifsy-1 regions were annotated as intact prophage via PHASTER (28.3% of *gogB*-harboring Dataset 1 [U.S. bovine and human data] genomes; Table 2, Supplementary Figure S19, and Supplementary Table S5). Occasionally, among Dataset 1 (U.S. bovine and human data) genomes, *gogB* was detected elsewhere in the genome: three genomes harbored *gogB* within regions annotated as prophage Gifsy-2 (3 of 180 *gogB*-harboring Dataset 1 [U.S. bovine and human data] genomes, 1.7%; Table 2 and Supplementary Table S5), while *gogB* was detected outside of annotated prophage regions within the remaining 51 *gogB*-harboring genomes (via PHASTER, 28.3% of *gogB*-harboring Dataset 1 [U.S. bovine and human data] genomes; Table 2 and Supplementary Table S5).

**Table 2.** Location of artAB and gogB in DT104 complex genomes within the four datasets queried in this study.

| Genes<sup>a</sup> | Dataset<br>(# of Genomes with Gene/<br>Total # of Genomes; %) | # of Genes Detected<sup>a</sup> |  |  |  |  |  | Outside of<br>Annotated<br>Prophage<br>Regions<br>(%)<sup>b,c</sup> |
| --- | --- | --- | --- | --- | --- | --- | --- | --- |
|  |  | Within Gifsy-1<sup>b</sup> |  | Within Gifsy-2<sup>b</sup> |  | Within *Salmonella*<br>phage 118970 sal3<sup>b</sup> |  |  |
|  |  | Intact<br>(%)<sup>c</sup> | Incomplete<br>(%)<sup>c</sup> | Intact<br>(%)<sup>c</sup> | Incomplete<br>(%)<sup>c</sup> | Intact<br>(%)<sup>c</sup> | Incomplete<br>(%)<sup>c</sup> |  |
| *artAB* | Dataset 1<sup>d</sup> (177/230; 77.0%) | 177<br>(100.0) | 0 (0) | 0 (0) | 0 (0) | 0 (0) | 0 (0) | 0 (0) |
|  | Dataset 2<sup>e</sup> (240/243; 98.8%) | 21 (8.8) | 0 (0) | 0 (0) | 0 (0) | 0 (0) | 0 (0) | 219 (91.3)<sup>h</sup> |
|  | Dataset 3<sup>f</sup> (271/290; 93.4%) | 263 (97.0) | 0 (0) | 0 (0) | 0 (0) | 2 (0.7) | 0 (0) | 6 (2.2)<sup>i</sup> |
|  | Dataset 4<sup>g</sup> (678/752; 90.2%) | 451 (66.5) | 0 (0) | 0 (0) | 0 (0) | 2 (0.3) | 0 (0) | 225 (33.2) |
| *gogB* | Dataset 1<sup>d</sup> (180/230; 78.3%) | 51 (28.3) | 75 (41.7) | 1 (0.6) | 2 (1.1) | 0 (0) | 0 (0) | 51 (28.3) |
|  | Dataset 2<sup>e</sup> (240/243; 98.8%) | 50 (20.8) | 85 (35.4) | 0 (0) | 0 (0) | 0 (0) | 0 (0) | 105 (43.8) |
|  | Dataset 3<sup>f</sup> (271/290; 93.4%) | 11 (4.1) | 67 (24.7) | 2 (0.7) | 1 (0.4) | 0 (0) | 0 (0) | 190 (70.1) |
|  | Dataset 4<sup>g</sup> (681/752; 90.6%) | 112 (16.4) | 223 (32.7) | 3 (0.4) | 3 (0.4) | 0 (0) | 0 (0) | 340 (49.9) |

<sup>a</sup> Identified using nucleotide BLAST (blastn; default settings, with no minimum identity or coverage threshold employed).
<sup>b</sup> Identified using the PHASTER webserver; “intact” refers to prophage classified by PHASTER as “intact”, while “incomplete” encompasses prophage classified as “incomplete” or “questionable”.
<sup>c</sup> Percentages in parentheses were calculated relative to the “# of genomes with gene” value in the “Dataset” column used as a denominator.
<sup>d</sup> Refers to the set of 230 U.S. human- and bovine-associated DT104 complex genomes identified and aggregated here.
<sup>e</sup> Refers to a set of 243 Scottish human- and bovine-associated DT104 genomes sequenced and characterized previously [102].
<sup>f</sup> Refers to a set of 290 DT104 genomes collected from various sources around the world, which were sequenced and characterized previously [13].
<sup>g</sup> Refers to the union of Dataset 1 (U.S. bovine and human data), Dataset 2 (Scottish bovine and human data), and Dataset 3 (multi-source data); eleven of the 290 genomes in Dataset 3 (multi-source data) were isolated from cattle and humans in the U.S. and thus were also included in Dataset 1 (U.S. bovine and human data).
<sup>h</sup> When 3 Kbp regions on either side of PHASTER prophage regions were considered, 234 Dataset 2 (Scottish bovine and human data) genomes harbored *artAB* within 3 Kbp of Gifsy-1 (97.5% of 240 *artAB*-harboring genomes in Dataset 2 [Scottish bovine and human data]); two and one genome harbored *artAB* within 3 Kb of other PHASTER prophage regions (annotated by PHASTER as Edward\_GF\_2\_NC\_026611 and PHAGE\_Enterо\_HK630\_NC\_019723, respectively).
<sup>i</sup> One genome (DK\_7322994\_6\_swine\_08-08-01) had *artA* and *artB* on separate contigs, with *artB* detected within 3 Kbp of an intact Gifsy-1 prophage region.

Only three genomes within Dataset 1 (U.S. bovine and human data) possessed an intact Gifsy-1 prophage via PHASTER but did not possess *artAB* (i.e., bovine-associated BOV_TYPH_Washington_2007_SRR1519881, BOV_TYPH_Minnesota_2010_SRR1089590, and BOV_TYPH_Minnesota_2008_SRR1177378, 1.7% of Dataset 1 [U.S. bovine and human data] genomes in which an intact Gifsy-1 was detected; Supplementary Figure S20 and Supplementary Tables S3 and S4). Interestingly, all three genomes possessed *gogB* (Supplementary Table S4). *gogB* was detected within an incomplete Gifsy-1 prophage region in the two genomes from Minnesota, while the genome from Washington did not harbor *gogB* within an annotated prophage region (via PHASTER; Tables 1 and 2, Supplementary Figure S20, and Supplementary Table S5).

Of the 168 bovine-associated Dataset 1 (U.S. bovine and human data) genomes, 150 (89.3%) possessed *artAB, gogB*, and Gifsy-1, while 153 (91.1%) possessed *gogB* and Gifsy-1 (Figure 3, Table 1, and Supplementary Figures S6-S8). Interestingly, of 62 human-associated Dataset 1 (U.S. bovine and human data) genomes, only 27 (43.5%) possessed *artAB, gogB,* and Gifsy-1 (Figure 3, Table 1, and Supplementary Figures S6-S8), indicating that Gifsy-1/*artAB*/*gogB* have a negative association with human-associated DT104 complex strains from the U.S. (two-sided FET raw *P*-value < 4.1×10^-12^, OR = 0.094; Table 1). However, no orthologous gene clusters within the Dataset 1 (U.S. bovine and human data) pan-genome shared a significant association with bovine or human host when accounting for population structure (treeWAS FDR-corrected *P*-value > 0.10).

Overall, 90 orthologous gene clusters within the Dataset 1 (U.S. bovine and human data) pan-genome were associated with Gifsy-1 presence or absence (via PHASTER; two-sided FET FDR-corrected *P*-value < 0.05, Supplementary Figure S13 and Supplementary Table S9). The presence and absence of 30 orthologous gene clusters shared a perfect association with Gifsy-1 presence and absence (via PHASTER; Supplementary Table S9). These genes were absent from all Dataset 1 (U.S. bovine and human data) genomes that did not possess Gifsy-1 and were present in all Dataset 1 (U.S. bovine and human data) genomes that did possess Gifsy-1 (FDR-corrected *P*-value < 0.05 and OR of infinity); in addition to *gogB*, these genes included numerous phage-associated proteins (Supplementary Table S9). Interestingly, genomes in which PHASTER did not detect an intact Gifsy-1 prophage region tended to possess a ColRNAI plasmid replicon (two-sided FET raw *P*-value < 1.0×10^-27^; Supplementary Figures S6-S8 and Supplementary Table S3).

### A MDR DT104 complex lineage circulating among cattle and humans across the U.S. lost *artAB*- and *gogB*-harboring prophage Gifsy-1 in the 1980s

To gain insight into the evolutionary relationships of *artAB*-negative U.S. DT104 complex strains, a time-scaled phylogeny was constructed using human- and bovine-associated U.S. DT104 complex genomes (i.e., genomes within Dataset 1 [U.S. bovine and human data]; Figure 3 and Supplementary Figures S6-S8). The common ancestor of the MDR U.S. bovine- and human-associated DT104 complex genomes included in this study was predicted to have existed circa 1975 (estimated node age 1974.9, node height 95% highest posterior density [HPD] interval [1958.1, 1986.4]; Figure 3 and Supplementary Figures S6-S8); this is consistent with observations made in previous studies [13, 114], in which DT104 was predicted to have acquired its MDR phenotype in the 1970s. The mean evolutionary rate estimated for the Dataset 1 (U.S. bovine and human data) genomes queried here was 1.75×10^-7^ substitutions/site/year (95% HPD interval [1.38×10^-7^, 2.11×10^-7^]), which is similar to evolutionary rates estimated in previous studies of DT104 isolates from other world regions [13, 102] (Supplementary Figure S5, Supplementary Table S8, and Supplementary Data).

Notably, over 75% of all Dataset 1 (U.S. bovine and human data) *artAB*-negative genomes (42 of 53 Dataset 1 [U.S. bovine and human data] *artAB*-negative genomes, 79.2%) were members of a single, well-supported clade (posterior probability = 1.0, referred to hereafter as the “U.S. *artAB*-negative major clade”; Figure 3, Supplementary Figures S6-S8, and Supplementary Table S15). In addition to lacking *artAB*, all members of the U.S. *artAB*-negative major clade lacked Gifsy-1 and 50 additional genes, which were present in over half of all Dataset 1 (U.S. bovine and human data) genomes not included in the U.S. *artAB*-negative major clade, including *gogB*, a chitinase, and many phage-associated proteins (Figure 3, Supplementary Figures S6-S8, and Supplementary Table S10).

Strains within the U.S. *artAB*-negative major clade were reportedly isolated between 1997 and 2018 (the most recent year included in this study) from at least 11 different states across the U.S. (for two isolates, the U.S. state in which the strain was isolated was unknown; Figures 2B and 3, Supplementary Figures S6-S8, and Supplementary Table S15). Interestingly, most strains within the U.S. *artAB*-negative major clade were isolated from humans (*n* = 30 of 42 U.S. *artAB*-negative major clade strains, 71.4%), and nearly half of all Dataset 1 (U.S. bovine and human data) genomes from human sources were members of this clade (*n* = 30 of 62 Dataset 1 [U.S. bovine and human data] genomes from human sources, 48.4%; Supplementary Tables S1 and S15). Human-associated U.S. *artAB*-negative major clade strains were reportedly isolated from six U.S. states between 1997 and 2014 (Supplementary Table S15). The majority of human-associated strains were isolated in Pennsylvania (*n* = 22 of 30 human-associated U.S. *artAB*-negative major clade strains, 73.3%; Figure 2B and Supplementary Table S15); however, Pennsylvania strains were reportedly isolated over a five-year period (i.e., from 2009 to 2014; Supplementary Table S15) and showcased considerable genomic diversity (Figure 3), indicating that it is highly unlikely that all human cases have an epidemiological link (i.e., they were not sequenced as part of a single, point-source outbreak). Bovine strains within the U.S. *artAB*-negative major clade were isolated from cattle or beef products (*n* = 12 of 42 U.S. *artAB*-negative major clade genomes, 28.6%; Figure 2B and Supplementary Table S15). Much like their human-associated counterparts, bovine-associated members of the U.S. *artAB*-negative major clade were interspersed throughout the clade’s phylogeny and varied in terms of isolation date (i.e., 2004 to 2018) and geographic origin (i.e., six U.S. states; Figure 2B and Supplementary Table S15).

Based on results of ancestral state reconstruction using *artAB* presence/absence, the loss of Gifsy-1, *artAB*, *gogB*, and other Gifsy-1-associated genes among members of the U.S. *artAB*-negative major clade was estimated to have occurred between 1985 and 1987 (estimated node ages 1985.0 and 1987.2, node height 95% HPD intervals [1979.0, 1990.2] and [1981.7, 1992.1], respectively; Figure 3 and Supplementary Figures S10-S12). This predicted loss event occurred circa a predicted rapid increase in the U.S. DT104 complex effective population size in the mid-to-late 1980s (Figure 4 and Supplementary Figure S9). Following this predicted rapid increase in the 1980s, the U.S. DT104 complex effective population size was predicted to have increased again in the mid-to-late 1990s, peaking circa 2000 (Figure 4 and Supplementary Figure S9).

### Loss of *artAB* and *gogB* within the global DT104 complex population occurs sporadically

The absence of Gifsy-1, *artAB*, and/or *gogB* among DT104 complex strains was not strictly a U.S. phenomenon: Gifsy-1, *artAB*, and *gogB* were not detected in three and 19 genomes out of (i) 243 DT104 strains isolated from cattle and humans in Scotland (referred to here as “Dataset 2 [Scottish bovine and human data]”) [102], and (ii) 290 DT104 strains collected from numerous sources around the world (referred to here as “Dataset 3 [multi-source data]”) [13], respectively (representing 1.2% and 6.6% of genomes in Dataset 2 [Scottish bovine and human data] and Dataset 3 [multi-source data], respectively; Figure 5, Table 1, Supplementary Figures S14 and S15, and Supplementary Tables S3-S5). Overall, out of 752 total DT104 complex genomes queried in this study (i.e., the union of Dataset 1 [U.S. bovine and human data], Dataset 2 [Scottish bovine and human data], and Dataset 3 [multi-source data], referred to here as “Dataset 4 [combined global dataset]”), *artAB* could not be detected in 74 genomes (9.8% of 752 Dataset 4 [combined global dataset] genomes; Table 1 and Supplementary Tables S15 and S16).

The Gifsy-1/*artAB*/*gogB* loss event associated with the U.S. *artAB*-negative major clade represented the single largest *artAB* loss event observed in this study (*n* = 42 of 752 total Dataset 4 [combined global dataset] genomes; Figure 5 and Supplementary Figures S14 and S15). However, several additional, sporadic *artAB* loss events among clades encompassing ≤5 genomes were observed (Figure 5, Supplementary Figures S14 and S15, and Supplementary Table S16). Overall, the 32 *artAB*-negative genomes that did not belong to the U.S. *artAB*-negative major clade were isolated from (i) a variety of sources (i.e., humans, cattle, pigs, and poultry), (ii) on four continents (i.e., North America, Europe, Asia, and Oceania), (iii) between 1992 and 2015 (Supplementary Table S16).

Among all 752 Dataset 4 (combined global dataset) genomes, the presence and absence of *artAB* and *gogB* was correlated with that of Gifsy-1 (two-sided FET raw *P*-value < 2.2×10^-16^ for each, ORs of 2069.8 and infinity, respectively), as well as each other (two-sided FET raw *P*-value < 2.2×10^-16^, infinite OR; Figure 5, Table 1, and Supplementary Figure S14). However, unlike the 177 *artAB*-harboring Dataset 1 (U.S. bovine and human data) genomes queried here, *artAB* was not always detected within prophage regions annotated as Gifsy-1 in other datasets (i.e., genomes in Dataset 2 [Scottish bovine and human data] and Dataset 3 [multi-source data]; Table 2, Supplementary Figure S21, and Supplementary Table S5). Two genomes from Dataset 3 (multi-source data) harbored *artAB* within regions annotated as *Salmonella* phage 118970_sal3 (using PHASTER’s nomenclature, “PHAGE_Salmon_118970_sal3_NC_031940”; Table 2 and Supplementary Table S5). However, despite not being annotated by PHASTER as “Gifsy-1”, these prophage regions shared a high degree of sequence homology with the DT104 Gifsy-1 prophage (Supplementary Figure S21).

Notably, for over 90% of *artAB*-harboring genomes in Dataset 2 (Scottish bovine and human data), *artAB* was identified outside of prophage regions annotated by PHASTER (*n* = 219 of 240 *artAB*-harboring genomes in Dataset 2 [Scottish bovine and human data]; Table 2 and Supplementary Table S5). However, when 3 Kbp regions on either side of PHASTER prophage regions were considered, 234 Dataset 2 (Scottish bovine and human data) genomes harbored *artAB* within 3 Kbp of Gifsy-1 (97.5% of 240 *artAB*-harboring genomes in Dataset 2 [Scottish bovine and human data]). Two and one genome harbored *artAB* within 3 Kb of other PHASTER prophage regions (annotated by PHASTER as Edward_GF_2_NC_026611 and PHAGE_Entero_HK630_NC_019723, respectively). For Dataset 3 (multi-source data), six *artAB*-positive genomes did not harbor *artAB* within PHASTER prophage regions (Table 2 and Supplementary Table S5). For one of these genomes (DK_7322994_6_swine_08-08-01), *artA* and *artB* were detected on separate contigs, with *artB* present within 3 Kbp of a PHASTER prophage region annotated as intact Gifsy-1; for the remaining five genomes, *artAB* was not located within a 5 Kb region upstream or downstream of any PHASTER prophages (Table 2 and Supplementary Table S5).

### *In vitro* response of U.S. DT104 complex strains to human- and bovine-associated gastrointestinal stress factors is not correlated with the presence of *artAB*- and *gogB*-harboring Gifsy-1

The (i) loss of Gifsy-1/*artAB*/*gogB* associated with the U.S. *artAB*-negative major clade circa a predicted rapid increase in the U.S. DT104 complex effective population size, plus (ii) the over-representation of human strains in the U.S. *artAB*-negative major clade led us to hypothesize that ArtAB and/or GogB production (or some other genomic element harbored on Gifsy-1) may influence the dynamics of DT104 complex strains in the digestive tracts of human and bovine hosts. Thus, we used phenotypic assays that simulated human and/or bovine digestion-associated stress conditions to compare the phenotypes of Gifsy-1/*artAB*/*gogB*-negative members of the U.S. *artAB*-negative major clade to those of the most closely related, Gifsy-1/*artAB*/*gogB*-positive U.S. DT104 complex strains available (Supplementary Table S11).

As the first three compartments of the bovine digestive tract differ massively from that of the human gut, the phenotype of Gifsy-1/*artAB*/*gogB-*positive and -negative strains was investigated in fresh bovine ruminal fluid (RF) obtained from a donor cow (Supplementary Table S12). DT104 complex concentrations were reduced by 3.4 log CFU (SD=0.2) when inoculated into RF at a final concentration of 10^5^ CFU/mL, whereas DT104 complex numbers were reduced by 1.3 log CFU (SD = 0.2) when inoculated at a final concentration of 10^8^ CFU/mL (Figure 6). While the inoculation density did significantly affect survival (ANOVA raw *P*-value < 0.001), the phenotype in RF was not associated with the presence or absence of Gifsy-1/*artAB*/*gogB* (ANOVA raw *P*-value > 0.05; Figure 6).

**Figure 6.**
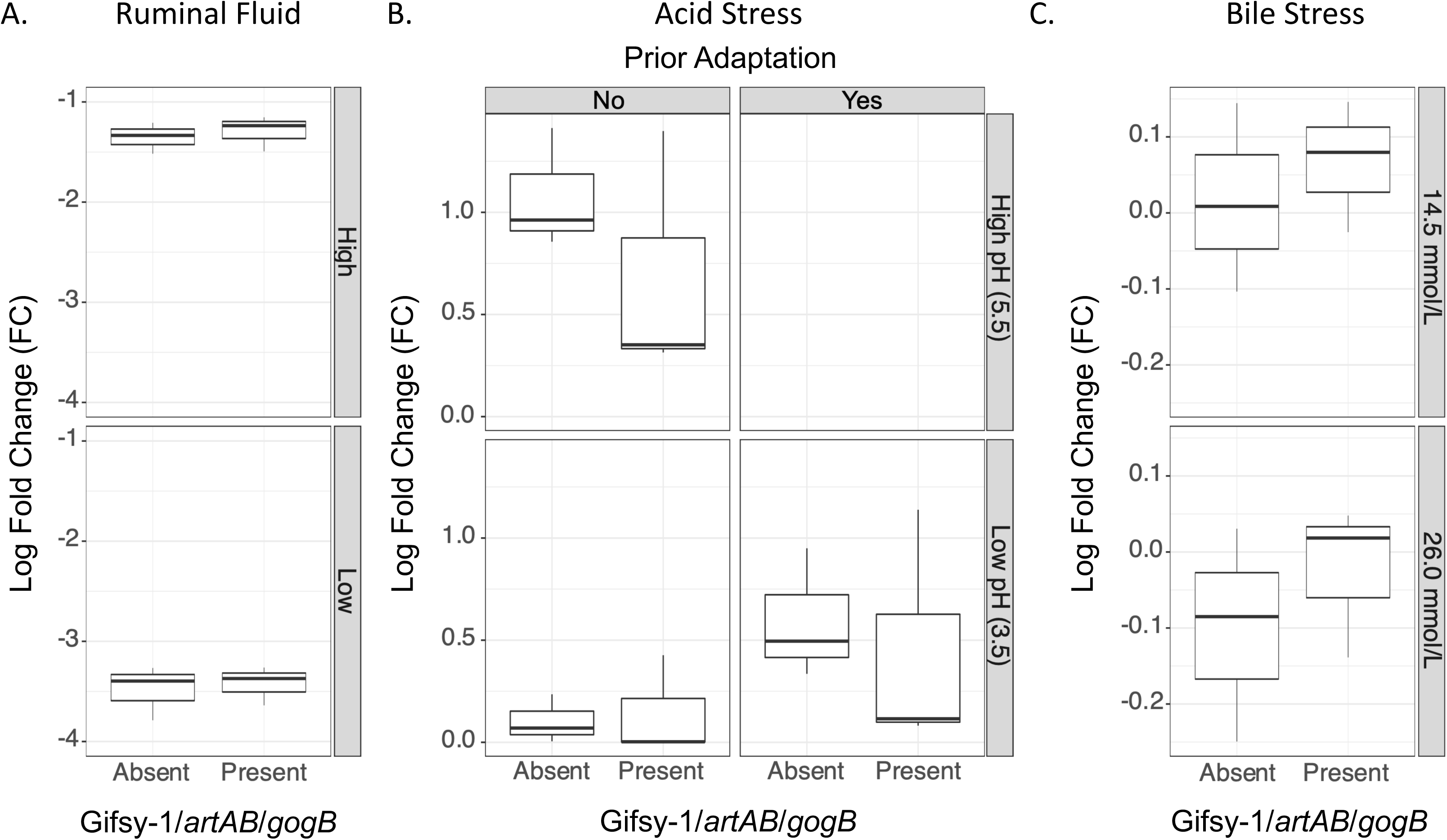
Response of DT104 complex isolates (*n* = 6) to environmental stress factors within the context of Gifsy-1/*artAB*/*gogB* presence and absence. Base-ten logarithmic fold change (FC) was calculated as follows: FC = log CFU/g at the start of the experiments – log CFU/g after the stress assay. (A) Log FC of DT104 inoculated into ruminal fluid at high (10^8^ CFU/mL; “High”) or low (10^5^ CFU/mL; “Low”) bacterial numbers. (B) Log FC of DT104 isolates exposed to inorganic acid stress (pH 3.5) with or without a prior adaption step with an intermediate pH (pH 5.5). (C) Log FC of DT104 isolates after exposure to bile salt at two concentrations.

The presence of Gifsy-1/*artAB*/*gogB* also did not significantly influence acid stress survival at pH 3.5 (ANOVA raw *P*-value > 0.05; Figure 6 and Supplementary Table S13). While prior adaptation at an intermediate pH 5.5 significantly increased survival at pH 3.5 as expected (ANOVA raw *P*-value < 0.01), there was no significant difference in acid adaptation between Gifsy-1/*artAB*/*gogB*-positive and -negative strains (ANOVA raw *P*-value > 0.05; Figure 6). Both groups showed a concentration-dependent reduction in growth/survival at the two tested bile concentrations of 0.6% and 1.1% (ANOVA raw *P* = 0.01 for the difference in fold change at the two concentrations), but there was no Gifsy-1/*artAB*/*gogB*-dependent phenotype in the response of DT104 complex strains to bile stress (ANOVA raw *P*-value > 0.05; Figure 6 and Supplementary Table S14).

## DISCUSSION

### Members of the DT104 complex largely harbor *artAB* on Gifsy-1-like prophage

Bacterial ADP-ribosylating toxins play important roles in the virulence of numerous pathogens [19, 115]. While the illness caused by *Salmonella enterica* is not considered to be a toxin-mediated disease in the classical sense (e.g., as is the case for *Clostridium botulinum* or *Vibrio cholerae*) [19], some *Salmonella* lineages are capable of producing ADP-ribosylating toxins, allowing them to alter host immune responses and promote pathogenesis [17, 19, 116–118]. ArtAB is one such toxin with a variable presence among *Salmonella* lineages: genes encoding ArtAB have been detected in at least 88 different serotypes and are correlated with the presence of typhoid toxin genes [24], although in DT104 this is not the case [17, 25]. Additionally, in the majority of these serotypes, *artA* is predicted to be a pseudogene and the selective advantage of maintaining *artB* appears to be related to its use as an alternative binding subunit for the typhoid toxin [19, 119].

A previous study of ArtAB-producing DT104 strains [18] found that ArtAB production among DT104 appears to be the norm rather than the exception, as 237 of 243 strains (97.5%) in the study were *artAB*-positive [18]. We observed similar findings here, as *artAB* was detected in 678 of 752 DT104 complex genomes (90.2%; Table 1 and Supplementary Table S4). Among U.S. human- and bovine-associated DT104 complex genomes (i.e., Dataset 1 [U.S. bovine and human data]), *artAB* was exclusively harbored on prophage regions identified by PHASTER as intact Gifsy-1 (177 of 177 *artAB*-harboring Dataset 1 [U.S. bovine and human data] genomes, 100%; Table 2). It’s important to note that Gifsy-1 prophage regions—as defined via PHASTER—can display a significant degree of genetic heterogeneity [28]. Furthermore, Gifsy-1 has been shown to share homology with Gifsy-2 [120, 121], a result observed here (Figure 1A and Supplementary Figures S1 and S17-S20). Thus, defining a “ground-truth” Gifsy-1 prophage in the genomic era may not necessarily be straightforward, as both Gifsy-1 and Gifsy-2 were originally identified in 1997 using Southern blotting [120]. Considering PHASTER results (Table 2 and Supplementary Table S5), our own sequence homology observations (Supplementary Figures S17-S20), and the fact that previous studies of DT104 have reportedly identified *artAB* within Gifsy-1 [10, 19], we are confident that the *artAB*-harboring prophage identified in Dataset 1 (U.S. bovine and human data) can be safely referred to as Gifsy-1.

Among genomes from other sources and/or world regions (i.e., Dataset 2 [Scottish bovine and human data] and Dataset 3 [multi-source data]), *artAB* was not always detected within the bounds of prophage regions annotated as Gifsy-1 (Table 2). Two *artAB*-harboring prophages that were not annotated as “Gifsy-1” by PHASTER, for example, shared a high degree of sequence homology with DT104 Gifsy-1 (Supplementary Figure S21), further highlighting the challenges associated with genomic differentiation of homology-sharing prophages defined in the pre-genomics era. However, most notably, *artAB* was frequently detected outside of annotated prophage regions in Dataset 2 (Scottish bovine and human data; Table 2). When 3 Kbp regions upstream and downstream of PHASTER prophage regions were considered, nearly all *artAB*-harboring Dataset 2 (Scottish bovine and human data) genomes possessed *artAB* within 3 Kbp of Gifsy-1. While it’s possible that *artAB* is indeed harbored outside of Gifsy-1 in these genomes, we suspect this is an artifact of prophage boundary prediction. Precise prediction of prophage boundaries is challenging [122], and additional factors (e.g., assembly fragmentation in prophage regions) can further affect boundary accuracy [45]. We thus encourage readers to interpret these results with caution. Future long-read sequencing efforts will thus likely provide much-needed insight into the prophage repertoire of the DT104 complex.

### A DT104 complex lineage isolated across multiple U.S. states for over twenty years lost its ability to produce the ArtAB toxin and anti-inflammatory effector GogB

Here, we observed that *artAB* loss events appear sporadically throughout the DT104 complex phylogeny (Figures 3 and 5). Among U.S. human- and bovine-associated DT104 complex genomes (i.e., Dataset 1 [U.S. bovine and human data]), these loss events usually coincided with Gifsy-1 loss, although not exclusively (i.e., three strains did not possess *artAB*, but possessed Gifsy-1; Figure 3 and Supplementary Figure S20). As mentioned above, *in silico* prophage detection and differentiation is challenging, and it is possible that HGT within a Gifsy-1-like prophage led to the loss of *artAB*, rather than the complete excision of Gifsy-1 in its entirety. In this scenario, it is plausible that partial Gifsy-1 remnants within these genomes would not be classified as Gifsy-1 prophage elements, or they would not be detected by *in silico* prophage detection methods. Regardless, we have identified 30 prophage-associated genes, which were detected in all Gifsy-1-harboring genomes and absent from all Gifsy-1-negative genomes in Dataset 1 (U.S. bovine and human data), indicating that numerous prophage-associated genes were lost along with *artAB* and *gogB* (Supplementary Table S9).

Most notably, we observed a MDR DT104 complex clade circulating among cattle and humans across 11 U.S. states, which lost Gifsy-1, concomitant with the ability to produce ArtAB and GogB (i.e., the U.S. *artAB*-negative major clade; Figure 3). Considering (i) genomic diversity observed within the U.S. *artAB*-negative major clade, along with the fact that (ii) U.S. *artAB*-negative major clade members have been isolated from multiple states and sources for over twenty years, it is nearly impossible that U.S. *artAB*-negative major clade genomes are the result of repeated sequencing of identical or nearly identical strains (e.g., as would be the case in a point-source outbreak scenario; Supplementary Table S15). When U.S. *artAB*-negative major clade genomes were compared to DT104 complex genomes collected from a variety of isolation sources around the world (Figure 5), we did not identify genomes from any country other than the U.S. within this clade, nor did we identify genomes from any isolation source other than cattle and humans (Figure 5 and Supplementary Table S15). Furthermore, we observed a high proportion of human-associated strains within the U.S. *artAB*-negative major clade relative to bovine-associated strains (Figure 3 and Supplementary Table S15). The (i) limited host range, (ii) limited geographic range, and (iii) high human-to-bovine ratio observed for the U.S. *artAB*-negative major clade in this study is likely an artifact of sampling and/or sequencing (e.g., due to the large number of bovine-associated genomes included in this study relative to other animal hosts, due to this study’s focus on the DT104 complex in the U.S., due to geographic and/or host biases in allocation of *Salmonella* sequencing resources). Future studies querying more DT104 complex strains from (i) non-human and non-bovine sources (e.g., other animal hosts, foods, environmental sources) and (ii) countries other than the U.S. and the United Kingdom will likely reveal a greater isolation source and geographic range for this clade, respectively.

However, it’s important to note that the U.S. *artAB*-negative major clade contained nearly half of all DT104 complex strains isolated from humans in the U.S. (*n* = 30 of 62 U.S. DT104 complex genomes from human sources, 48.4%; Supplementary Tables S1 and S15). This is notable, as our study included all human-associated U.S. DT104 complex genomes with metadata available in Enterobase at the time. It is certainly likely that there were U.S. DT104 complex genomes from human sources, which were not included in our study (e.g., due to missing publicly available metadata), and it is possible that there may be biases in terms of metadata reporting (e.g., some laboratories may routinely provide detailed, publicly available metadata for genomes that they sequence, while other laboratories may never or rarely provide metadata). Thus, in order to gain further insight into potential U.S. *artAB*-negative major clade host associations (or the lack thereof), it is essential that isolation source metadata are made publicly available in addition to WGS data.

The U.S. *artAB*-negative major clade was predicted to have lost Gifsy-1/*artAB*/*gogB* circa 1985-1987, circa a predicted rapid increase in the U.S. DT104 complex effective population size, which occurred in the mid-to-late 1980s (Figure 4). Our results are consistent with a previous study of DT104 from multiple world regions, which also identified periods of dramatic population growth in the 1980s and 1990s [13]. This rapid increase in population size is notable, as it coincides with the global MDR DT104 epidemic, which occurred among humans and animals throughout the 1990s [13, 14, 102]. However, it is essential to note that our data do not imply that *artAB*, *gogB*, or Gifsy-1 loss played a role in the emergence and subsequent global spread of DT104; any potential association between the virulence and/or fitness of MDR DT104 and Gifsy-1/*artAB*/*gogB* loss among DT104 complex genomes is merely speculative at this point. While previous studies of DT104 have shown that prophage excision and *artAB* loss occur in response to DNA damage and other stressors [17, 19], future studies are needed to better understand the roles that Gifsy-1, *artAB*, and *gogB* play in DT104 evolution.

### Members of the U.S. *artAB*-negative major clade do not have a phenotypic advantage relative to other U.S. DT104 complex strains when exposed to ruminal fluid-, acid-, and bile-associated stressors *in vitro*

*Salmonella enterica* encounters numerous stressors within the gastrointestinal tracts of humans and animals, including (but not limited to) low pH, low oxygen, exposure to bile, and the host immune system [123–125]. Furthermore, the gastrointestinal environment that *Salmonella enterica* encounters can differ between hosts; for example, the first three compartments of the bovine digestive tract differ massively from those of the human gut, as they essentially serve as massive microbial fermentation chambers [126]. Here, we evaluated the survival of DT104 complex strains when exposed to three stressors encountered in the human and/or bovine gastrointestinal tracts: (i) ruminal fluid (RF; bovine rumen), (ii) low pH (bovine abomasum and human stomach), and (iii) exposure to bile (bovine and human duodenum); we discuss each step in detail below.

In the bovine digestion process, the RF, including the complex community of ruminal microbiota [127], presents an early line of defense against potential pathogens, like *Salmonella* spp. In RF, the kill rate of DT104 complex strains was dependent on the inoculation density. The high inoculation rate (10^8^ CFU/mL) was chosen to test the ability of the ruminal microbiota to efficiently kill or impede *Salmonella*. The lower inoculation rate of 10^5^ CFU/mL was chosen for its dynamic range to measure either growth or decrease of *Salmonella* concentration. An interaction of the complex ruminal microbiota with the inoculated *Salmonella* is conceivable in two ways: either the microbiota exhibit strategies to produce antimicrobial compounds against *Salmonella* species [128, 129], or through competition for nutrients, e.g., iron [130]. The fact that the ruminal microbiota was less effective at killing DT104 complex strains at the high inoculation rate suggests that their defense mechanisms against DT104 complex strains are limited and/or the system started to be overrun by the high numbers of the DT104 complex strain.

Gastric acids in the stomach (or abomasum) are the next line of host defense, which *Salmonella* must overcome during gastrointestinal passage [131]. A pH of 3.5 was selected based on the following considerations: the human gastric pH varies from pH <2 in a fasted state to pH >6 during meals, returning to a low pH within hours postprandially [132, 133]. Intracellular pathogens like *Salmonella* spp. have adapted to survive low pH intracellularly in the phagosomes of phagocytes (e.g. pH 4-6) [134, 135] and express adaptive acid tolerance that allows them to tolerate pH values between 2-3 [136]. Therefore, the chosen pH 3.5 reflects a relevant physiological state of the human stomach and represents a sublethal stress to *Salmonella* spp. Our experiments confirmed that acid adaptation with HCl at pH 5.5 led to much higher survival rates at pH 3.5. Well-known mechanisms such as decreased membrane conductivity for H^+^, increased proton extrusion, or changes in the cell envelope composition [136–138] could be responsible for this.

Upon leaving the stomach, enteric pathogens are confronted with bile. Bile salts show antimicrobial activity by dissolving membrane lipids, dissociating integral membrane proteins [139], and lead to general cell damage by misfolding and denaturation of proteins [140, 141] and DNA damage [142, 143]. *Salmonella enterica* is able to survive duodenal bile salt concentrations through DNA repair mechanisms [143], multiple changes in gene expression [144], and increased production of anti-oxidative enzymes [145]. Here, selected DT104 complex strains were able to survive at both tested bile salt concentrations (14.5 mmol/L and 26.0 mmol/L); however, no significant differences were observed between strains that harbored Gifsy-1/*artAB*/*gogB* and those that did not (Figure 6).

In summary, the *in vitro* stress assays performed in this study aimed to mimic the stressors that DT104 complex strains encounter in the gastrointestinal tracts of humans and ruminants. Given the over-representation of human-associated Gifsy-1/*artAB*/*gogB*-negative strains observed here, one may be tempted to speculate that Gifsy-1, *artAB*, and/or *gogB* absence may confer members of the U.S. *artAB*-negative major clade with a competitive advantage in the human host gastrointestinal tract; however, no Gifsy-1/*artAB*/*gogB*-dependent phenotype was observed in DT104 complex strains under the tested conditions (Figure 6). Furthermore, as mentioned above, the overrepresentation of human strains in this clade could merely be an artifact of sampling/sequencing. Thus, it may be possible that Gifsy-1/*artAB*/*gogB* absence may confer some advantage(s) to U.S. *artAB*-negative major clade strains in hosts underrepresented in this study, or in environmental conditions, which were not tested in this study, including those outside of the host (e.g., high osmotic pressure and competitive microbiota in manure or wastewater, food safety measures like disinfectants, antimicrobials and food processing) [124]. However, at the present, this is merely speculation; future studies are needed to evaluate whether Gifsy-1/*artAB*/*gogB* loss among members of the U.S. *artAB*-negative major clade is merely coincidental or indicative of some evolutionarily advantageous phenotype.

### Future research is needed to understand the roles that Gifsy-1, ArtAB, and GogB play in DT104 virulence

The results presented here indicate that prophage-mediated ArtAB production within the DT104 complex can undergo temporal changes. Most notably, we identified the U.S. *artAB*-negative major clade, which lost the ability to produce ArtAB and GogB, likely due to a Gifsy-1 loss event (Figure 3). However, the ecological and/or evolutionary significance of this loss-of-function event remain unclear. Although phenotypic assessments have demonstrated a role for DT104-encoded ArtAB in both cell culture and a mouse model [18], the true benefit of this toxin in the context of human and bovine salmonellosis has not been investigated. It has been previously shown that reactive oxygen species (ROS) induce production of ArtAB [23], which may suggest that *artAB* is expressed in response to immune cell derived ROS. Furthermore, as treatment with ArtA increases intracellular levels of cAMP in macrophage-like cells [18], ArtAB may play a role in delaying *Salmonella* clearance by altering the activity of host immune cells [19]. Hence, future studies, including in tissue culture and animal models, will be needed to determine whether *artAB* presence or absence confers a selective advantage among human- and animal-associated DT104.

## AUTHOR STATEMENTS

### Conflicts of interest

The authors declare that there are no conflicts of interest.

### Funding information

This material is based on work supported by the National Science Foundation (NSF) Graduate Research Fellowship Program under grant no. DGE-1650441, with additional funding provided by an NSF Graduate Research Opportunities Worldwide (GROW) grant through a partnership with the Swiss National Science Foundation (SNF). This work was additionally supported by the SciLifeLab & Wallenberg Data Driven Life Science Program (grant: KAW 2020.0239).

## Acknowledgments

Figure 2 was created with BioRender.com.

## Notes

https://doi.org/10.5281/zenodo.7688792

### Competing Interest Statement

The authors have declared no competing interest.

### Summary of Updates

We have amended the text of the manuscript for clarity, and we have added additional Figures/Supplementary Material.

