## Supplementary Figure S1 for "A multidrug-resistant *Salmonella enterica* Typhimurium DT104 complex lineage circulating among humans and cattle in the United States lost the ability to produce pertussis-like toxin ArtAB"

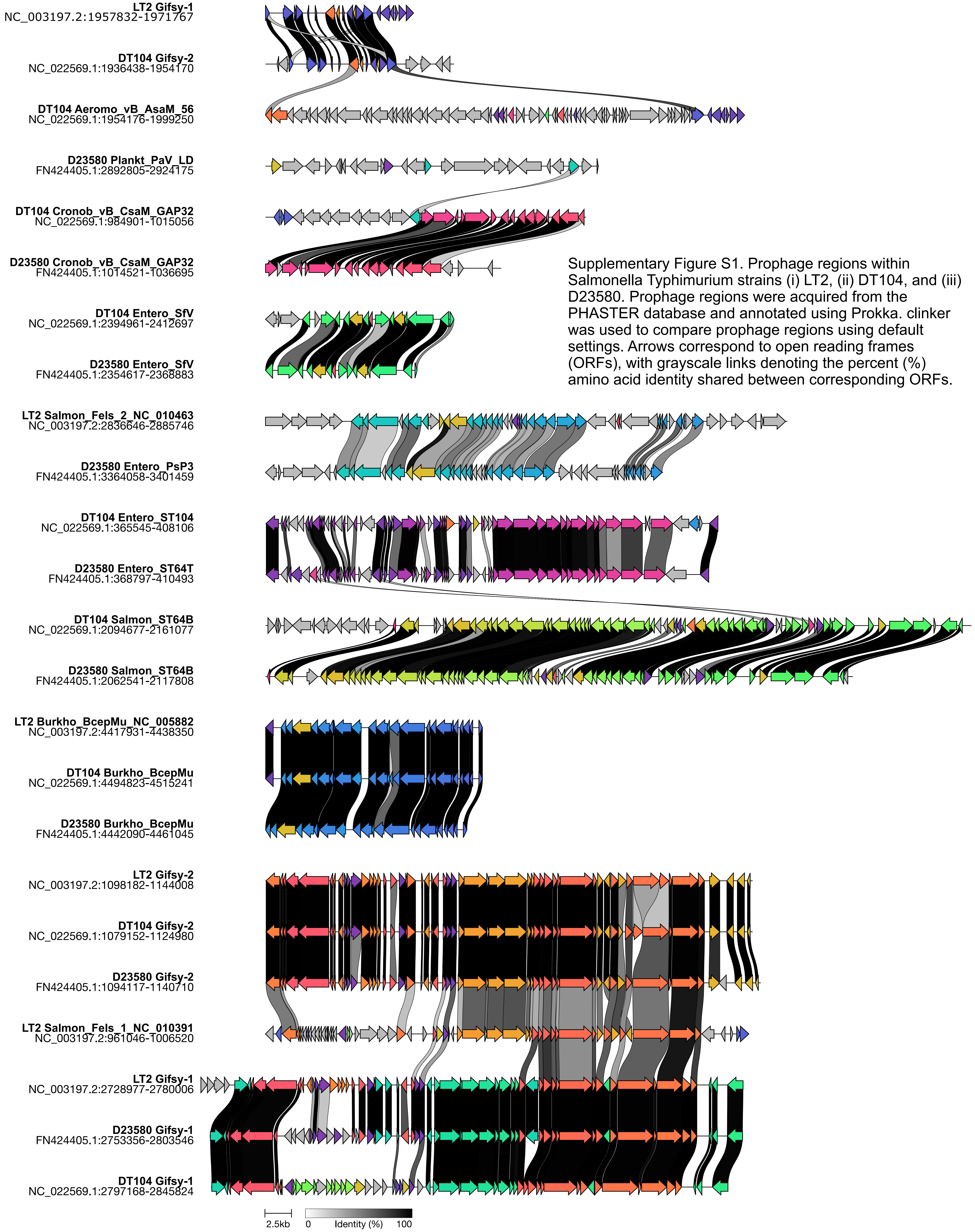


Supplementary Figure S1. Prophage regions within Salmonella Typhimurium strains (i) LT2, (ii) DT104, and (iii) D23580. Prophage regions were acquired from the PHASTER database and annotated using Prokka. clinker was used to compare prophage regions using default settings. Arrows correspond to open reading frames (ORFs), with grayscale links denoting the percent (%) amino acid identity shared between corresponding ORFs.
