## Supplementary Figure S2 for "A multidrug-resistant *Salmonella enterica* Typhimurium DT104 complex lineage circulating among humans and cattle in the United States lost the ability to produce pertussis-like toxin ArtAB"

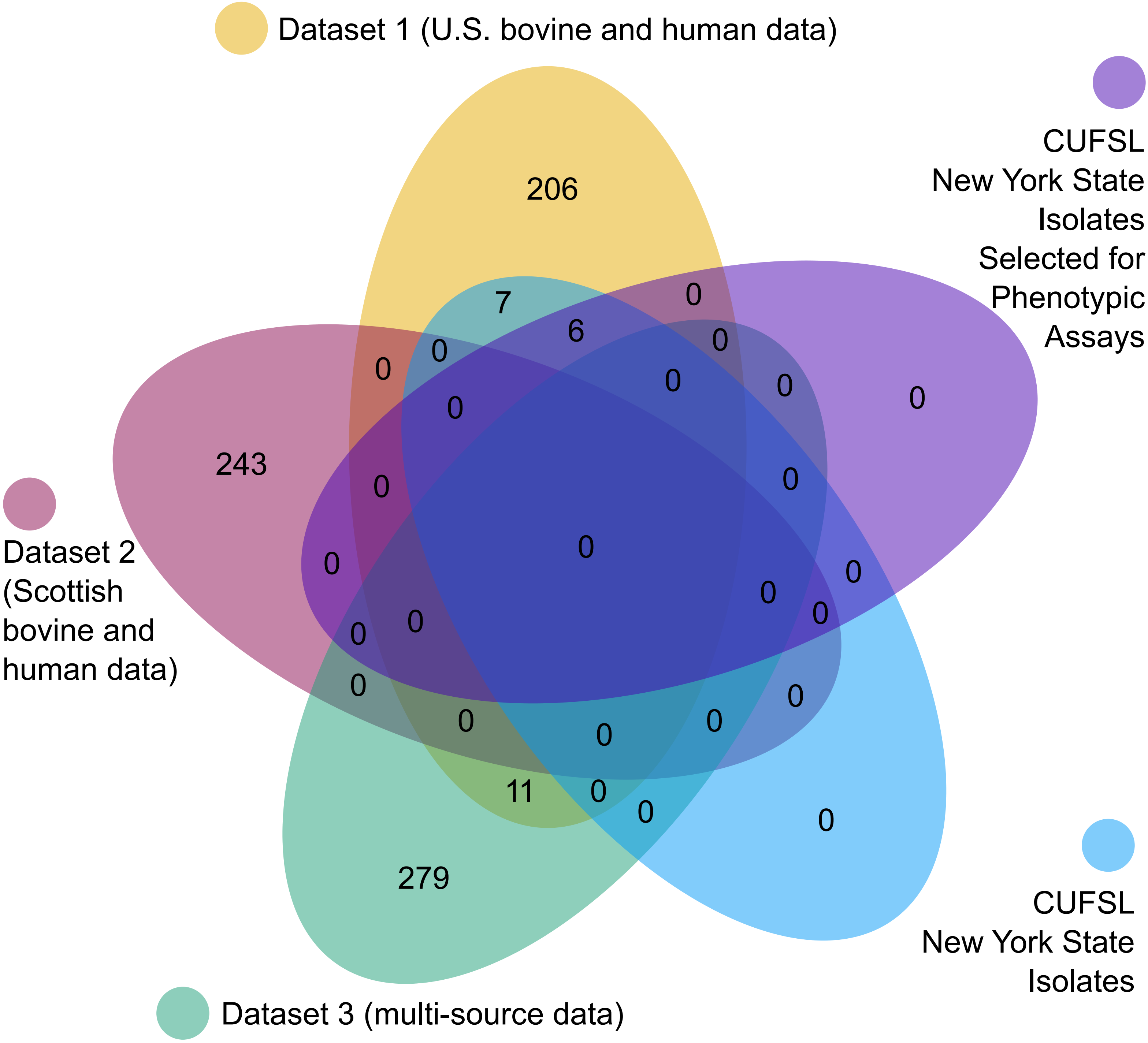


Supplementary Figure S2. Venn diagram showcasing the relationship between DT104 complex datasets used in this study. Numbers denote the number of genomes within a given dataset or subset of a dataset. For a flow chart with detailed descriptions of the datasets used in this study, see Supplementary Figure S3. CUFSL, Cornell University Food Safety Laboratory culture collection.
