## Supplementary Figure S3 for "A multidrug-resistant *Salmonella enterica* Typhimurium DT104 complex lineage circulating among humans and cattle in the United States lost the ability to produce pertussis-like toxin ArtAB"

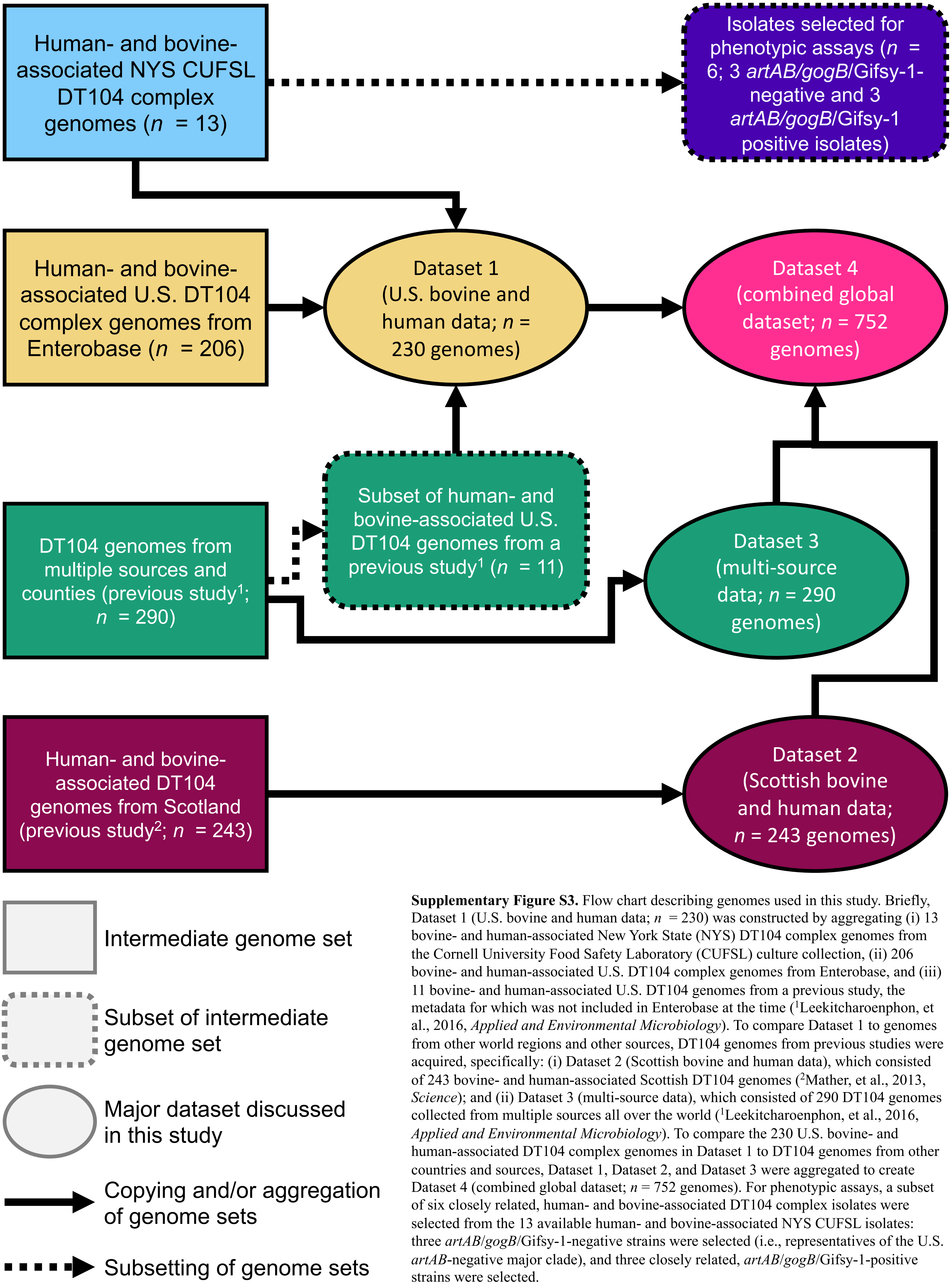


**Supplementary Figure S3.** Flow chart describing genomes used in this study. Briefly, Dataset 1 (U.S. bovine and human data; *n* = 230) was constructed by aggregating (i) 13 bovine- and human-associated New York State (NYS) DT104 complex genomes from the Cornell University Food Safety Laboratory (CUFSL) culture collection, (ii) 206 bovine- and human-associated U.S. DT104 complex genomes from Enterobase, and (iii) 11 bovine- and human-associated U.S. DT104 genomes from a previous study, the metadata for which was not included in Enterobase at the time ([1]Leekitcharoenphon, et al., 2016, *Applied and Environmental Microbiology*). To compare Dataset 1 to genomes from other world regions and other sources, DT104 genomes from previous studies were acquired, specifically: (i) Dataset 2 (Scottish bovine and human data), which consisted of 243 bovine- and human-associated Scottish DT104 genomes ([2]Mather, et al., 2013, *Science*); and (ii) Dataset 3 (multi-source data), which consisted of 290 DT104 genomes collected from multiple sources all over the world ([1]Leekitcharoenphon, et al., 2016, *Applied and Environmental Microbiology*). To compare the 230 U.S. bovine- and human-associated DT104 complex genomes in Dataset 1 to DT104 genomes from other countries and sources, Dataset 1, Dataset 2, and Dataset 3 were aggregated to create Dataset 4 (combined global dataset; *n* = 752 genomes). For phenotypic assays, a subset of six closely related, human- and bovine-associated DT104 complex isolates were selected from the 13 available human- and bovine-associated NYS CUFSL isolates: three *artAB*/*gogB*/Gifsy-1-negative strains were selected (i.e., representatives of the U.S. *artAB*-negative major clade), and three closely related, *artAB*/*gogB*/Gifsy-1-positive strains were selected.
