## Supplementary Figure S4 for "A multidrug-resistant *Salmonella enterica* Typhimurium DT104 complex lineage circulating among humans and cattle in the United States lost the ability to produce pertussis-like toxin ArtAB"

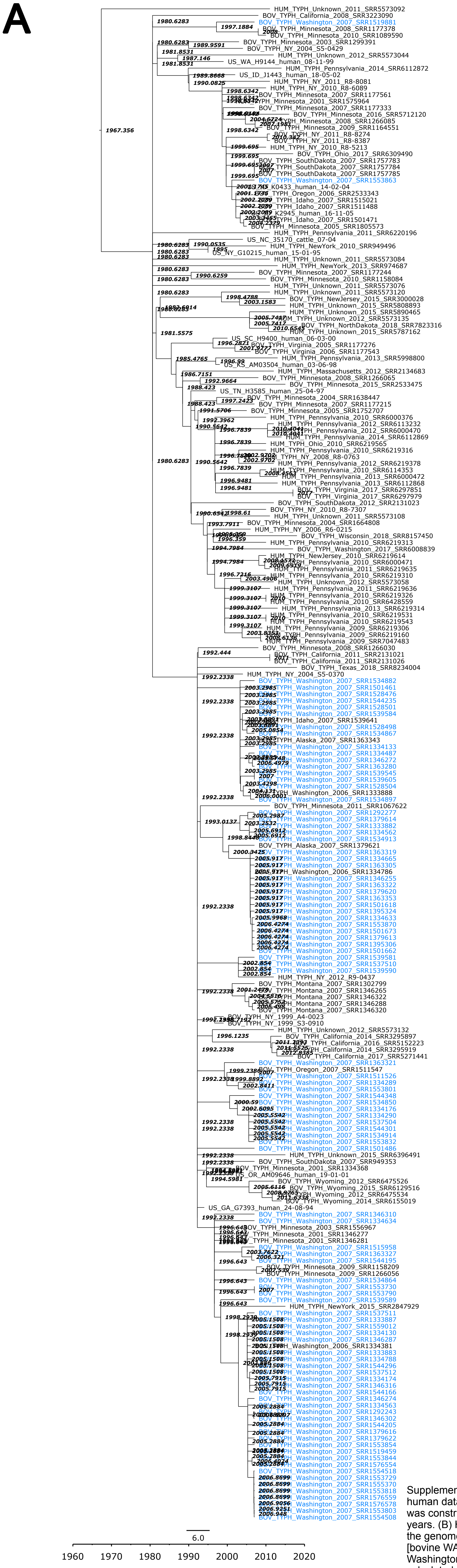


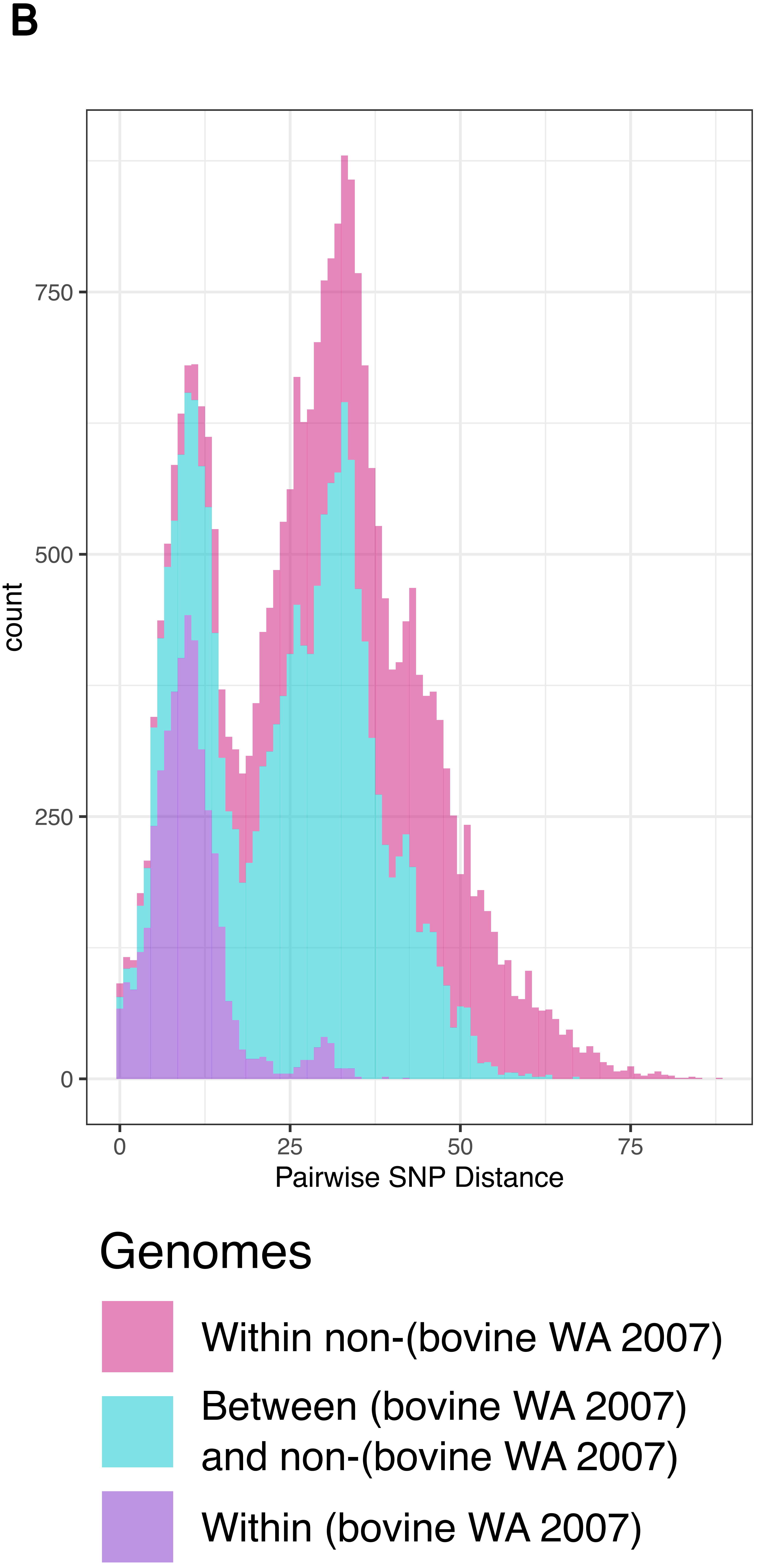


Supplementary Figure S4. (A) Time-scaled maximum likelihood (ML) phylogeny constructed using all 230 DT104 complex genomes in Dataset 1 (U.S. bovine and human data). Blue tip labels denote strains reportedly isolated from cattle in Washington State in 2007. Node labels correspond to node ages (year). The phylogeny was constructed using IQ-TREE, using core SNPs identified via Snippy as input. LSD2 was used to root and time-scale the phylogeny. Branch lengths are reported in years. (B) Histogram of pairwise SNP distances calculated between all bovine-associated U.S. DT104 complex genomes isolated in Washington State in 2007 (i.e., the genomes denoted by blue tip labels in panel A). Colored shading denotes distances calculated between (i) two 2007 bovine Washington State genomes (Within [bovine WA 2007]), (ii) two genomes not isolated in 2007 from bovine sources in Washington State (Within non-[bovine WA 2007]), and (iii) one 2007 bovine Washington State genome and one genome that was not a member of this set (Between [bovine WA 2007] and non-[bovine WA 2007]). Pairwise distances were calculated using the “dist.gene” function in the ape version 5.6-2 R package. The histogram was plotted using ggplot2.
