## Supplementary Figure S5 for "A multidrug-resistant *Salmonella enterica* Typhimurium DT104 complex lineage circulating among humans and cattle in the United States lost the ability to produce pertussis-like toxin ArtAB"

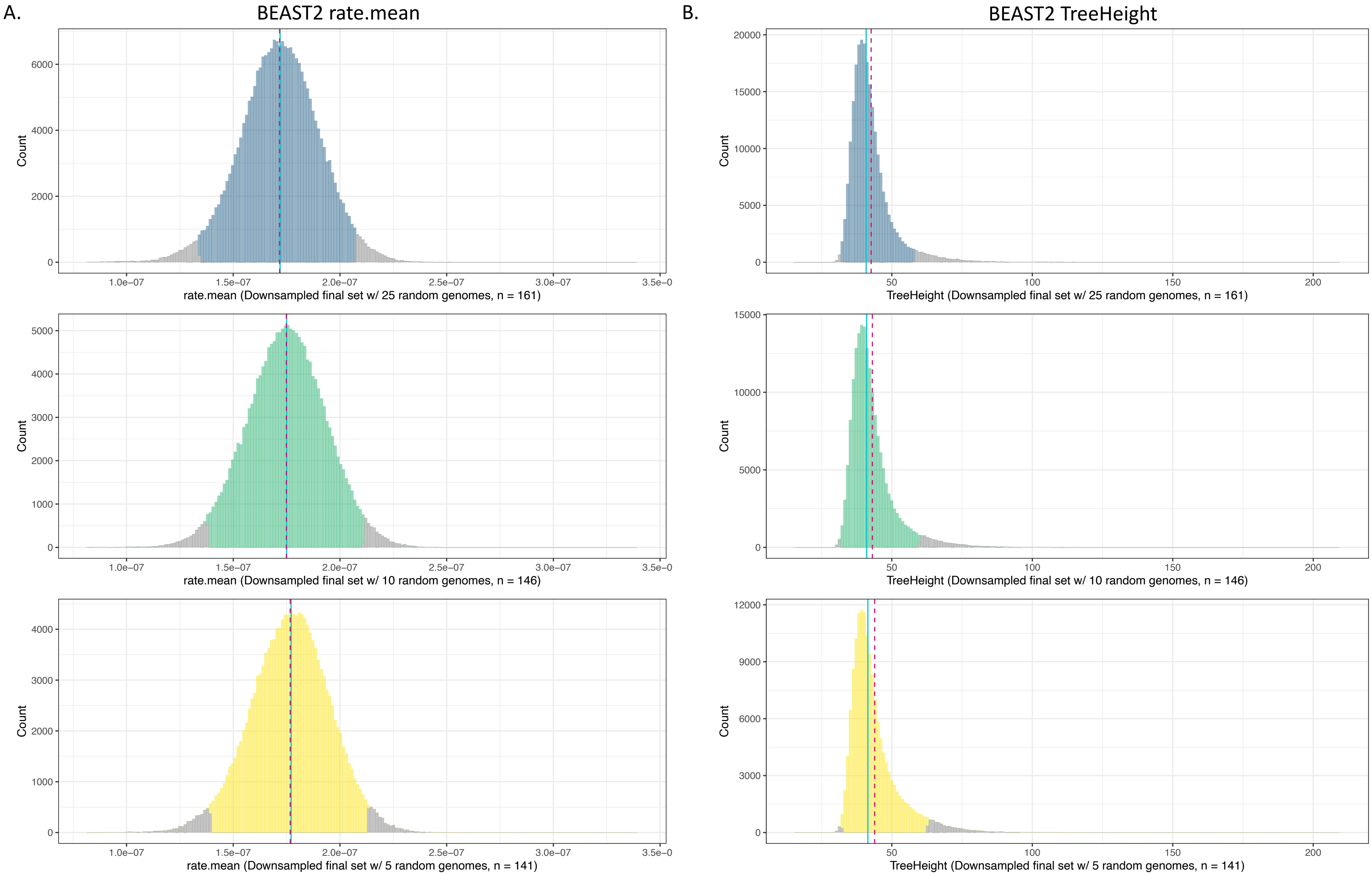


**Supplementary Figure S5.** Histograms of BEAST2 parameter estimates for the (A) rate.mean and (B) TreeHeight parameters for three downsampled Dataset 1 (U.S. bovine and human data) genome sets. The three genome sets were constructed by randomly selecting (i) 25, (ii) 10, and (iii) 5 bovine DT104 complex genomes collected in Washington State in 2007 from the complete set of 230 Dataset 1 (U.S. bovine and human data) genomes (*n* = 161, 146, and 141 total genomes in each downsampled genome set, respectively).
