## Supplementary Figure S6 for "A multidrug-resistant *Salmonella enterica* Typhimurium DT104 complex lineage circulating among humans and cattle in the United States lost the ability to produce pertussis-like toxin ArtAB"

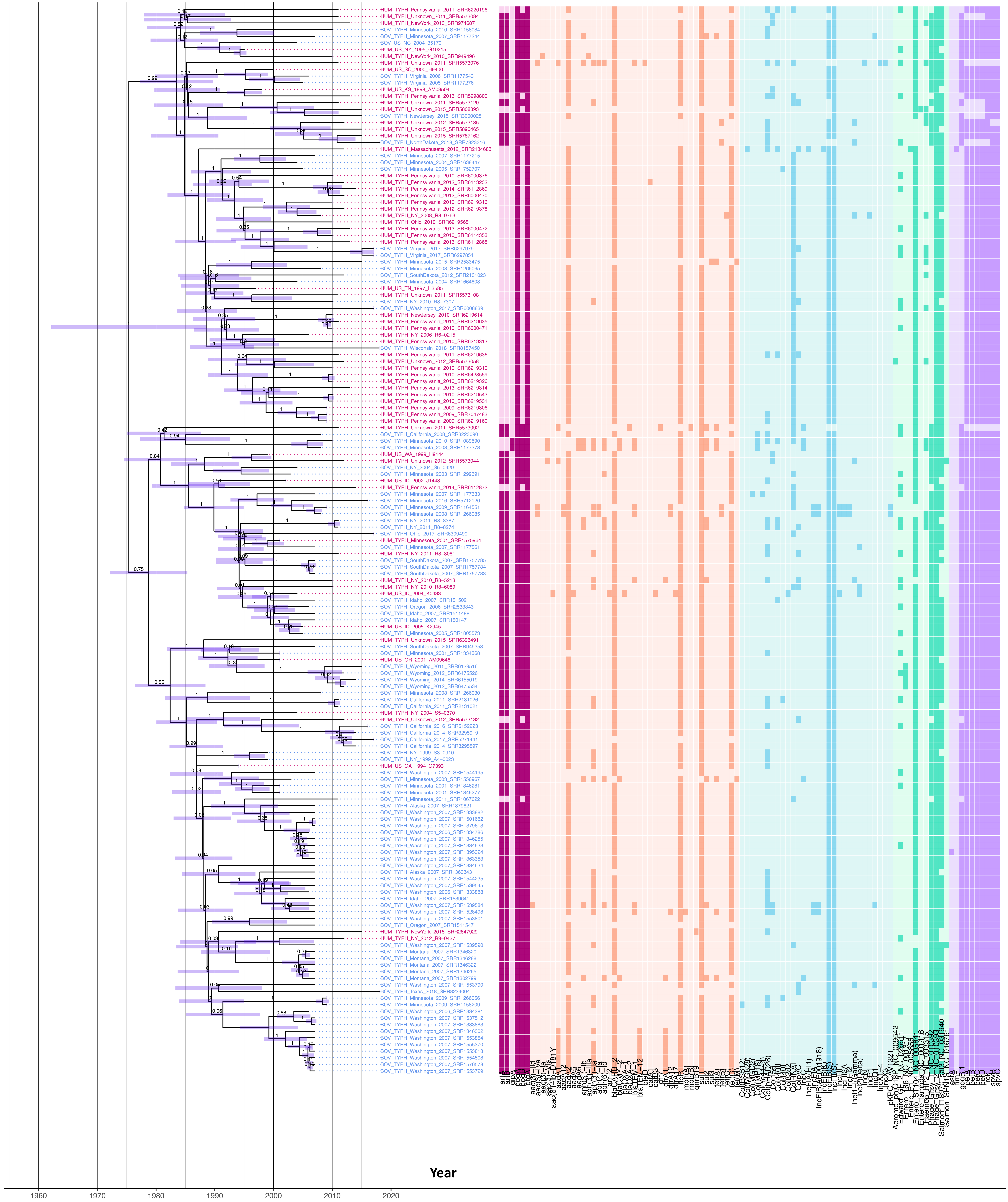


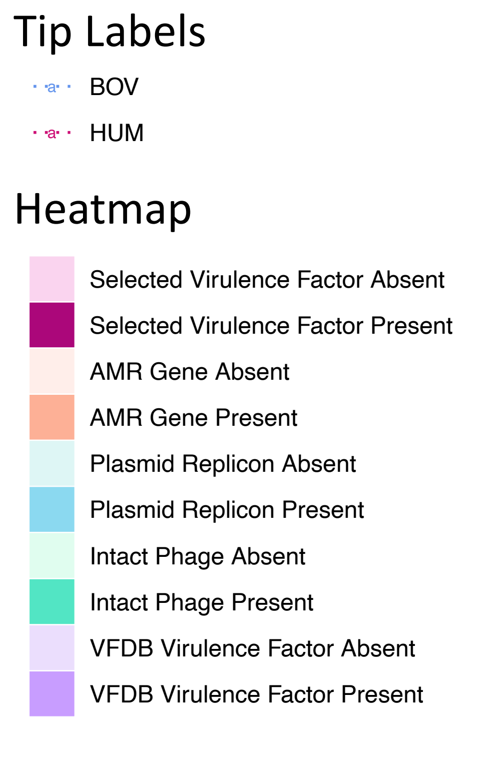


**Supplementary Figure S6.** Bayesian time-scaled phylogeny constructed using 161 human- and bovine-associated DT104 complex genomes collected in the United States (U.S.; i.e., the genome set downsampled from Dataset 1 [U.S. bovine and human data], using 25 randomly selected DT104 complex genomes collected from cattle in Washington State in 2007). Tip label colors denote the isolation source reported for each genome (human or bovine in pink and blue, respectively). The heatmap to the right of the phylogeny denotes the presence and absence of: (i) selected virulence factors (dark and light pink, respectively; selected virulence factors were detected using nucleotide BLAST and were considered present using a minimum coverage threshold of 40%); (ii) antimicrobial resistance (AMR) genes (dark and light orange, respectively; detected using ABRicate and the NCBI AMR database); (iii) plasmid replicons (dark and light blue, respectively; detected using ABRicate and the PlasmidFinder database); (iv) intact prophage (dark and light green, respectively; identified and classified as "intact" via PHASTER); (v) variably present Virulence Factor Database (VFDB) virulence factors (dark and light purple, respectively; detected using ABRicate and VFDB, with virulence factors detected in all 161 genomes omitted for readability). All analyses that relied on ABRicate employed minimum nucleotide identity and coverage thresholds of 75 and 50%, respectively. The phylogeny was constructed and rooted using BEAST2. Time in years is plotted along the X-axis, while branch labels correspond to posterior probabilities of branch support. Transparent purple node bars denote node height 95% highest posterior density (HPD) intervals.
