## Supplementary Figure S9 for "A multidrug-resistant *Salmonella enterica* Typhimurium DT104 complex lineage circulating among humans and cattle in the United States lost the ability to produce pertussis-like toxin ArtAB"

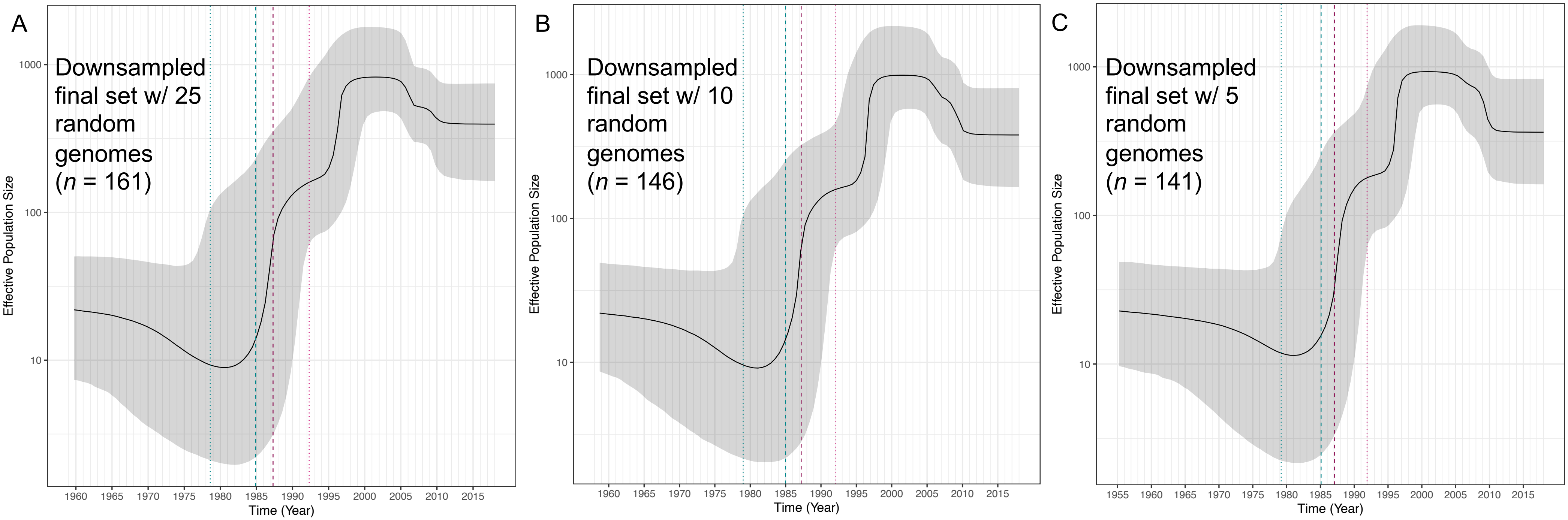


**Supplementary Figure S9.** Coalescent Bayesian Skyline plots constructed using Dataset 1 (U.S. bovine and human data) genomes, downsampled to (A) 25, (B) 10, and (C) 5 bovine genomes collected from Washington State in 2007 ($n$ = 161, 146, and 141 genomes, respectively). Effective population size and time in years are plotted on the Y- and X-axes, respectively. The median effective population size estimate is denoted by the solid black line, with upper and lower 95% highest posterior density (HPD) interval bounds denoted by gray shading. The interval bounded by dashed vertical lines denotes the time interval in which Gifsy-1/*artAB*/*gogB* were predicted to have been lost among members of the U.S. *artAB*-negative major clade. The dotted vertical lines correspond to the 95% HPD interval for Gifsy-1/*artAB*/*gogB* loss among members of the U.S. *artAB*-negative major clade.
