## Supplementary Figure S10 for "A multidrug-resistant *Salmonella enterica* Typhimurium DT104 complex lineage circulating among humans and cattle in the United States lost the ability to produce pertussis-like toxin ArtAB"

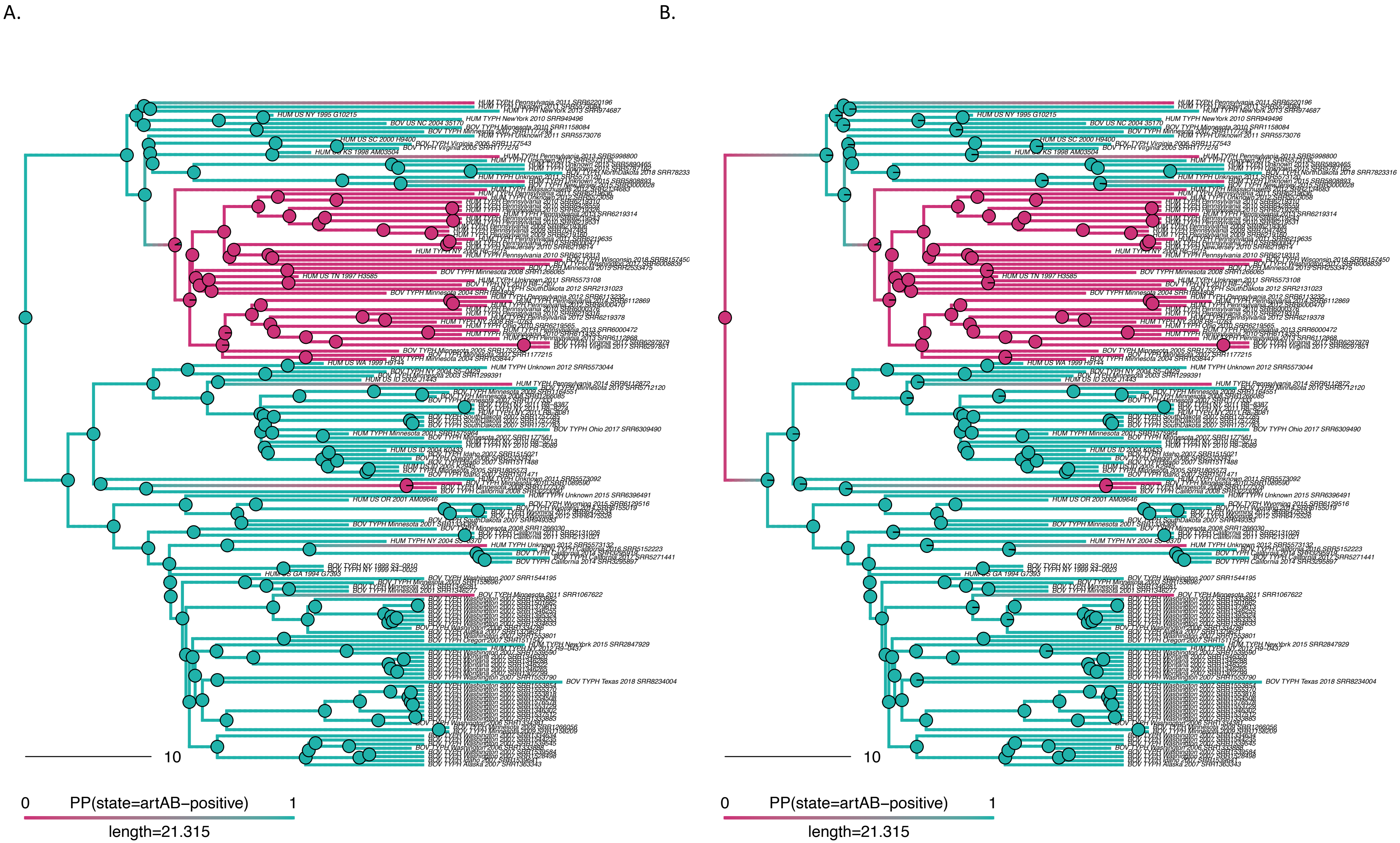


**Supplementary Figure S10.** Time-scaled Bayesian phylogeny of 161 human- and bovine-associated U.S. DT104 genomes (i.e., the genome set downsampled from Dataset 1 [U.S. bovine and human data], using 25 randomly selected DT104 complex genomes collected from cattle in Washington State in 2007). Tree edge and node colors correspond to the posterior probability (PP) of being in an *artAB*-positive state, obtained using an empirical Bayes approach, in which a continuous-time reversible Markov model was fitted, followed by 10,000 simulations of stochastic character histories using the fitted model and tree tip states. Root node prior probabilities for *artAB*-positive and *artAB*-negative states were (A) equal (i.e., 0.5 each) or (B) estimated using the make.simmap function in the phytools package in R. Branch lengths are reported in years.
