## Supplementary Figure S11 for "A multidrug-resistant *Salmonella enterica* Typhimurium DT104 complex lineage circulating among humans and cattle in the United States lost the ability to produce pertussis-like toxin ArtAB"

A. B.

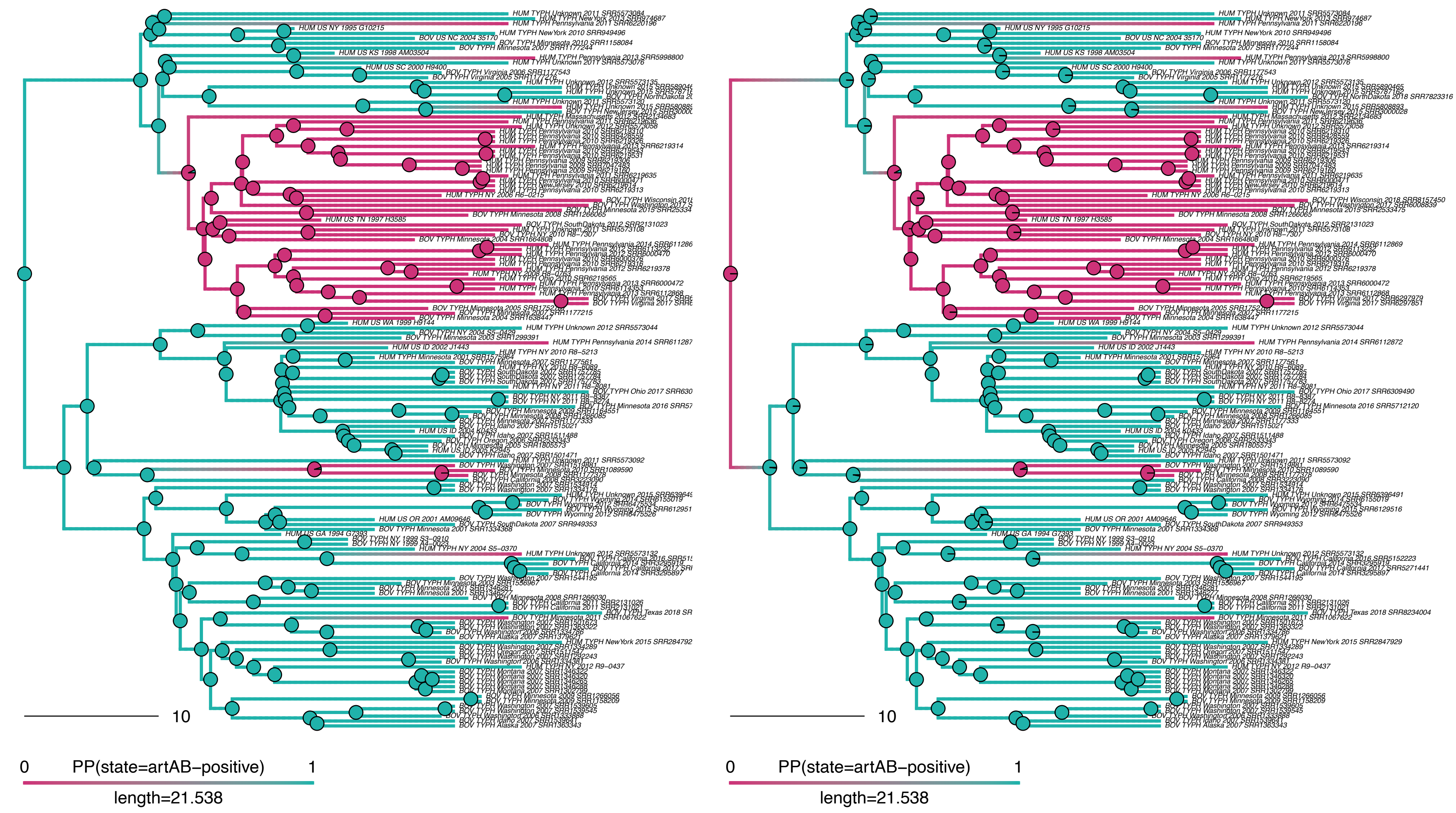


**Supplementary Figure S11.** Time-scaled Bayesian phylogeny of 146 human- and bovine-associated U.S. DT104 complex genomes (i.e., the genome set downsampled from Dataset 1 (U.S. bovine and human data), using 10 randomly selected DT104 complex genomes collected from cattle in Washington State in 2007). Tree edge and node colors correspond to the posterior probability (PP) of being in an *artAB*-positive state, obtained using an empirical Bayes approach, in which a continuous-time reversible Markov model was fitted, followed by 10,000 simulations of stochastic character histories using the fitted model and tree tip states. Root node prior probabilities for *artAB*-positive and *artAB*-negative states were (A) equal (i.e., 0.5 each) or (B) estimated using the make.simmap function in the phytools package in R. Branch lengths are reported in years.
