## Supplementary Figure S13 for "A multidrug-resistant *Salmonella enterica* Typhimurium DT104 complex lineage circulating among humans and cattle in the United States lost the ability to produce pertussis-like toxin ArtAB"

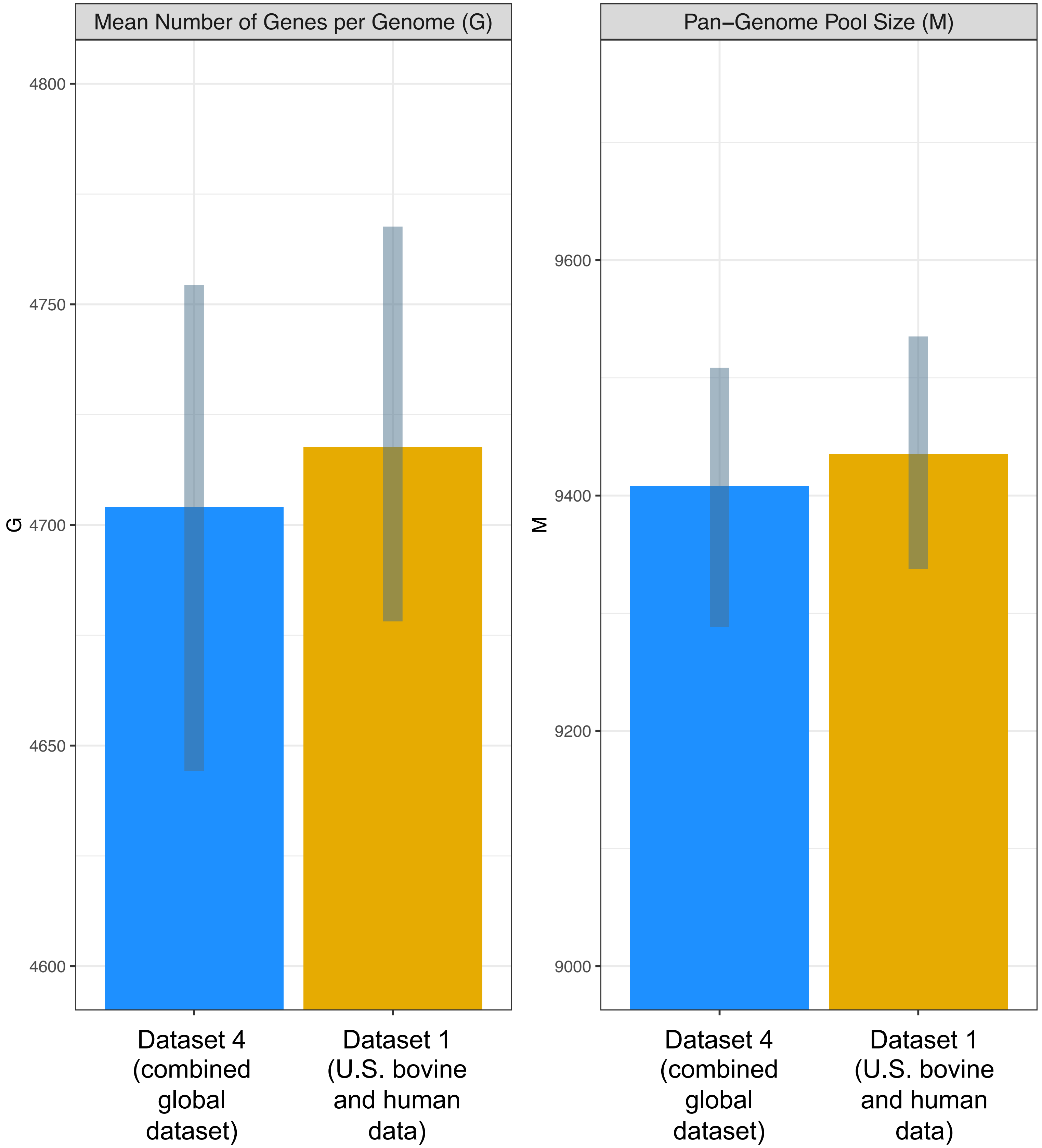


**Supplementary Figure S13.** Inferred parameters for the Finite Many Genes (FMG) model for the following data sets: (i) Dataset 4 (combined global dataset) ($n$ = 752 DT104 complex genomes); (ii) Dataset 1 (U.S. bovine and human data) ($n$ = 230 DT104 complex genomes). FMG parameters were estimated using Panaroo, with gray bars denoting the 2.5 and 97.5% confidence interval bounds for each parameter (obtained using 100 bootstrap replicates).
