## Supplementary Figure S14 for "A multidrug-resistant *Salmonella enterica* Typhimurium DT104 complex lineage circulating among humans and cattle in the United States lost the ability to produce pertussis-like toxin ArtAB"

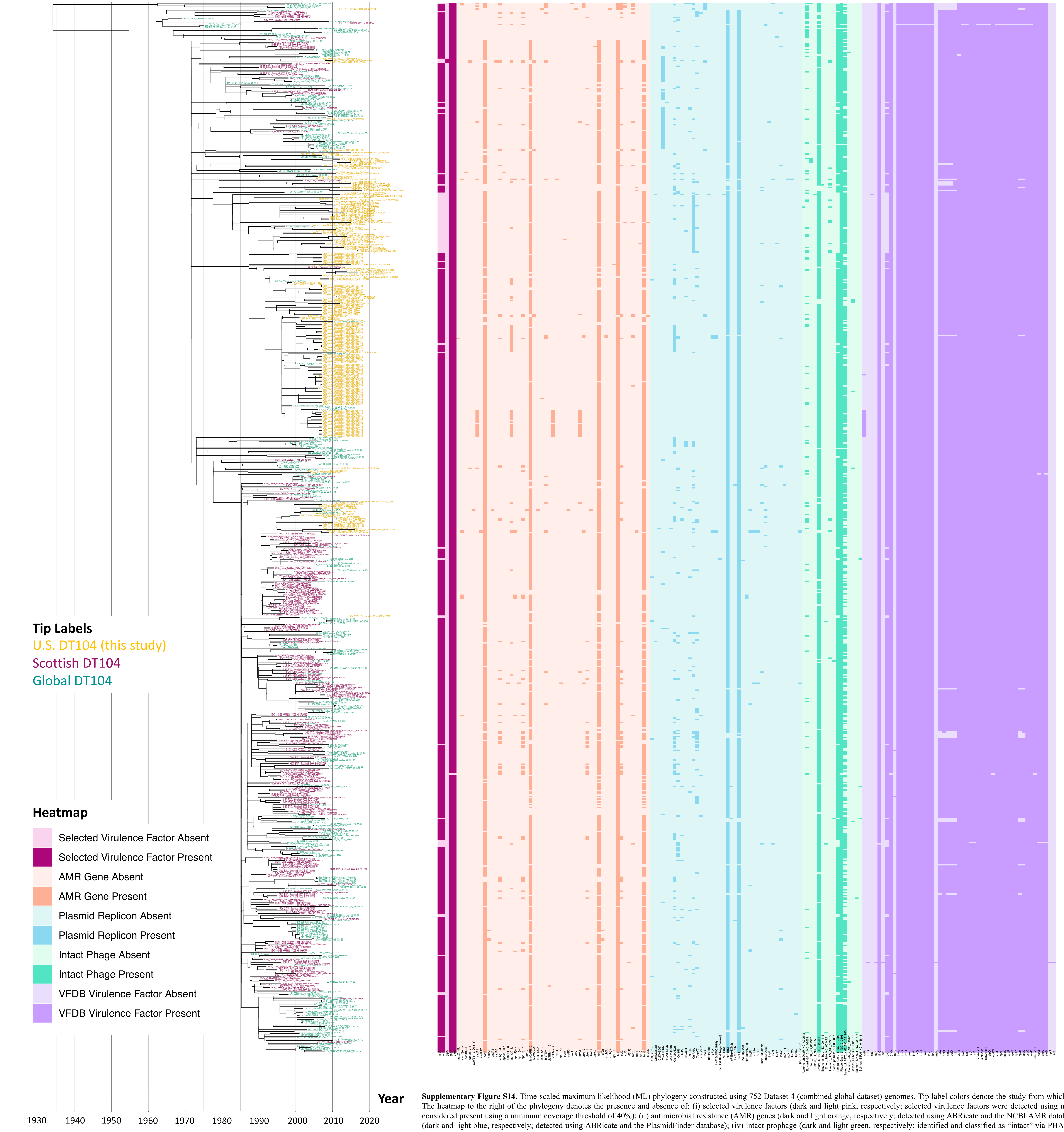


**Supplementary Figure S14.** Time-scaled maximum likelihood (ML) phylogeny constructed using 752 Dataset 4 (combined global dataset) genomes. Tip label colors denote the study from which each genome was derived. The heatmap to the right of the phylogeny denotes the presence and absence of: (i) selected virulence factors (dark and light pink, respectively; selected virulence factors were detected using nucleotide BLAST and were considered present using a minimum coverage threshold of 40%); (ii) antimicrobial resistance (AMR) genes (dark and light orange, respectively; detected using ABRicate and the NCBI AMR database); (iii) plasmid replicons (dark and light blue, respectively; detected using ABRicate and the PlasmidFinder database); (iv) intact prophage (dark and light green, respectively; identified and classified as "intact" via PHASTER); (v) variably present Virulence Factor Database (VFDB) virulence factors (dark and light purple, respectively; detected using ABRicate and VFDB, with virulence factors detected in all 752 genomes omitted for readability). All analyses that relied on ABRicate employed minimum nucleotide identity and coverage thresholds of 75 and 50%, respectively. The phylogeny was constructed using IQ-TREE and rooted and time-scaled using LSD2. Time in years is plotted along the X-axis.
