## Supplementary Figure S16 for "A multidrug-resistant *Salmonella enterica* Typhimurium DT104 complex lineage circulating among humans and cattle in the United States lost the ability to produce pertussis-like toxin ArtAB"

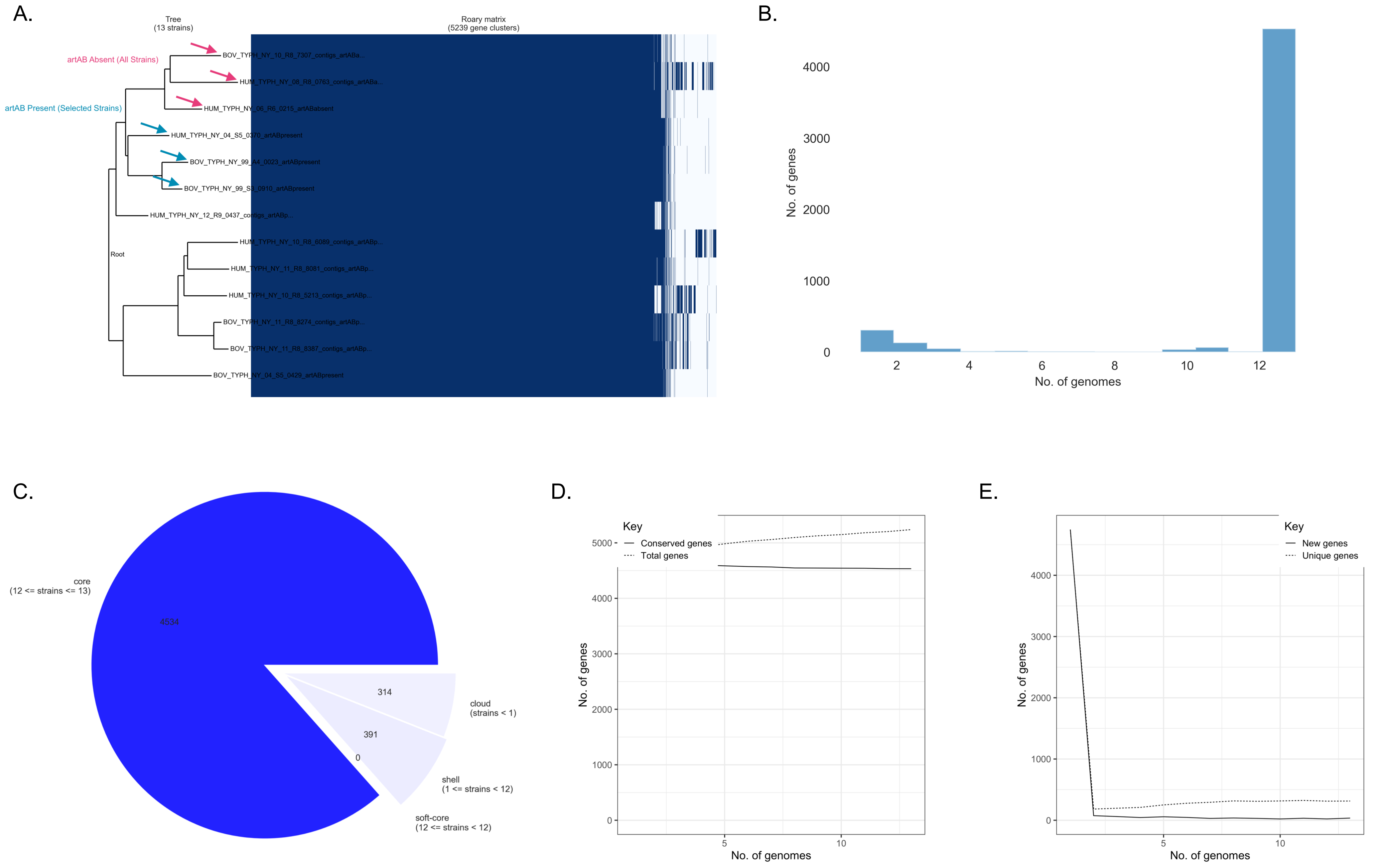


**Supplementary Figure S16.** (Pan-)genomic characterization of the 13 Cornell University Food Safety Laboratory (CUFSL) DT104 complex strains, which were available for phenotypic assays. (A) Roary gene presence/absence matrix. The phylogeny to the left of the matrix corresponds to the maximum likelihood (ML) phylogeny produced via Parsnp, rooted at the midpoint. *artAB*-positive and *artAB*-negative strains selected to undergo phenotypic characterization are denoted by blue and pink arrows, respectively. (B) Histogram of the number of genes detected among the 13 genomes. (C) Pie chart showcasing pan-/core-genome composition. (D) Number of conserved and total and (E) new and unique genes among the 13 DT104 complex genomes. Plots (A-C) were constructed using roary_plots.py version 0.1.0 (https://github.com/sanger-pathogens/Roary/blob/master/contrib/roary_plots/roary_plots.py). Plots D and E were produced via Roary.
