## Supplementary Figure S19 for "A multidrug-resistant *Salmonella enterica* Typhimurium DT104 complex lineage circulating among humans and cattle in the United States lost the ability to produce pertussis-like toxin ArtAB"

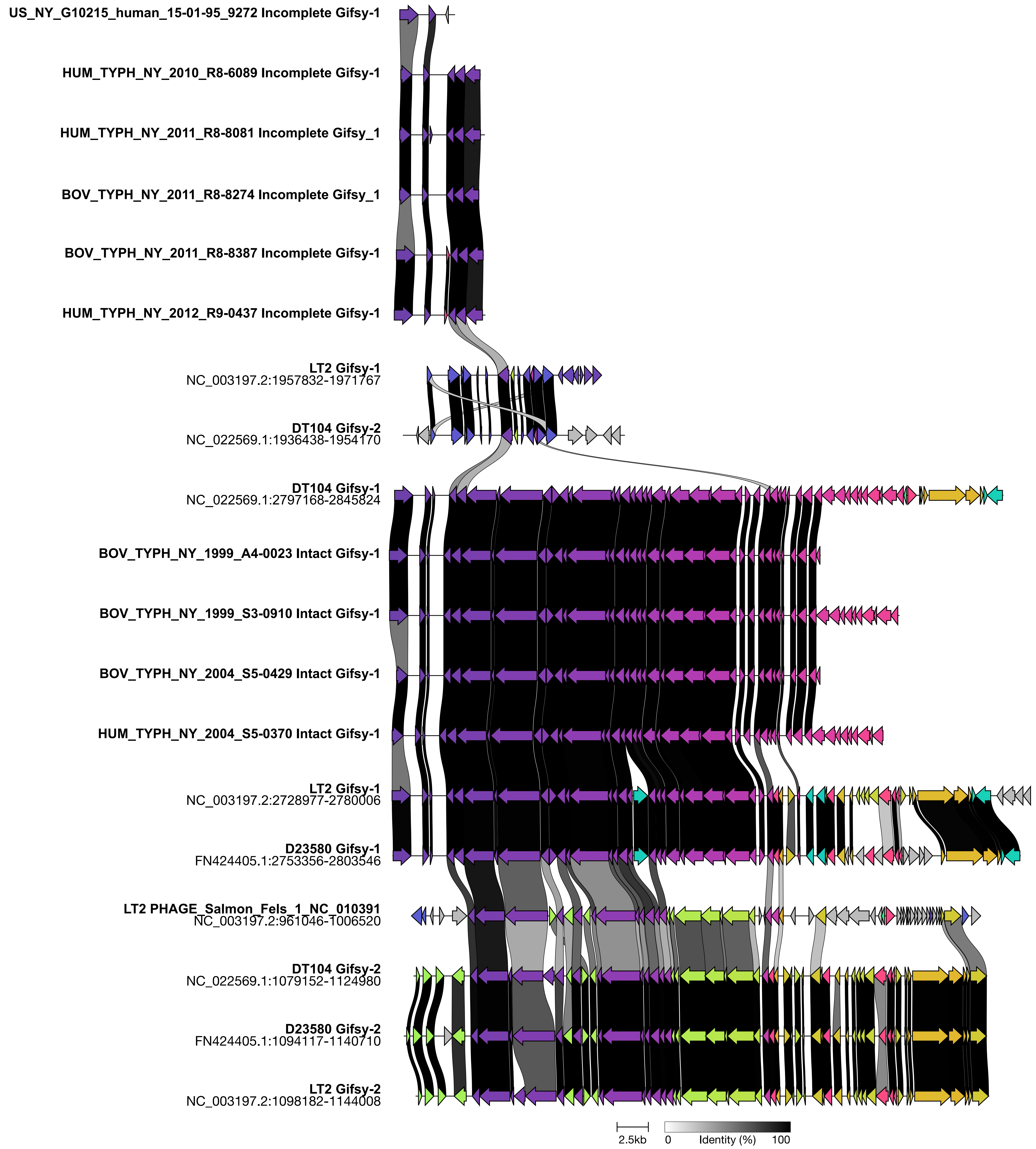


Supplementary Figure S19. gogB-harboring prophage regions in human- and bovine-associated DT104 complex genomes from New York State, as compared to Gifsy-1- and Gifsy-2-like prophages described in Salmonella Typhimurium strains (i) LT2, (ii) DT104, and (iii) D23580. gogB-harboring prophage regions were detected in the New York State genomes using PHASTER. Selected LT2, DT104, and D23580 prophage regions were acquired from the PHASTER database. All prophage regions were annotated using Prokka. clinker was used to compare prophage regions using default settings. Arrows correspond to open reading frames (ORFs), with grayscale links denoting the percent (%) amino acid identity shared between corresponding ORFs.
