## Supplementary Figure S20 for "A multidrug-resistant *Salmonella enterica* Typhimurium DT104 complex lineage circulating among humans and cattle in the United States lost the ability to produce pertussis-like toxin ArtAB"

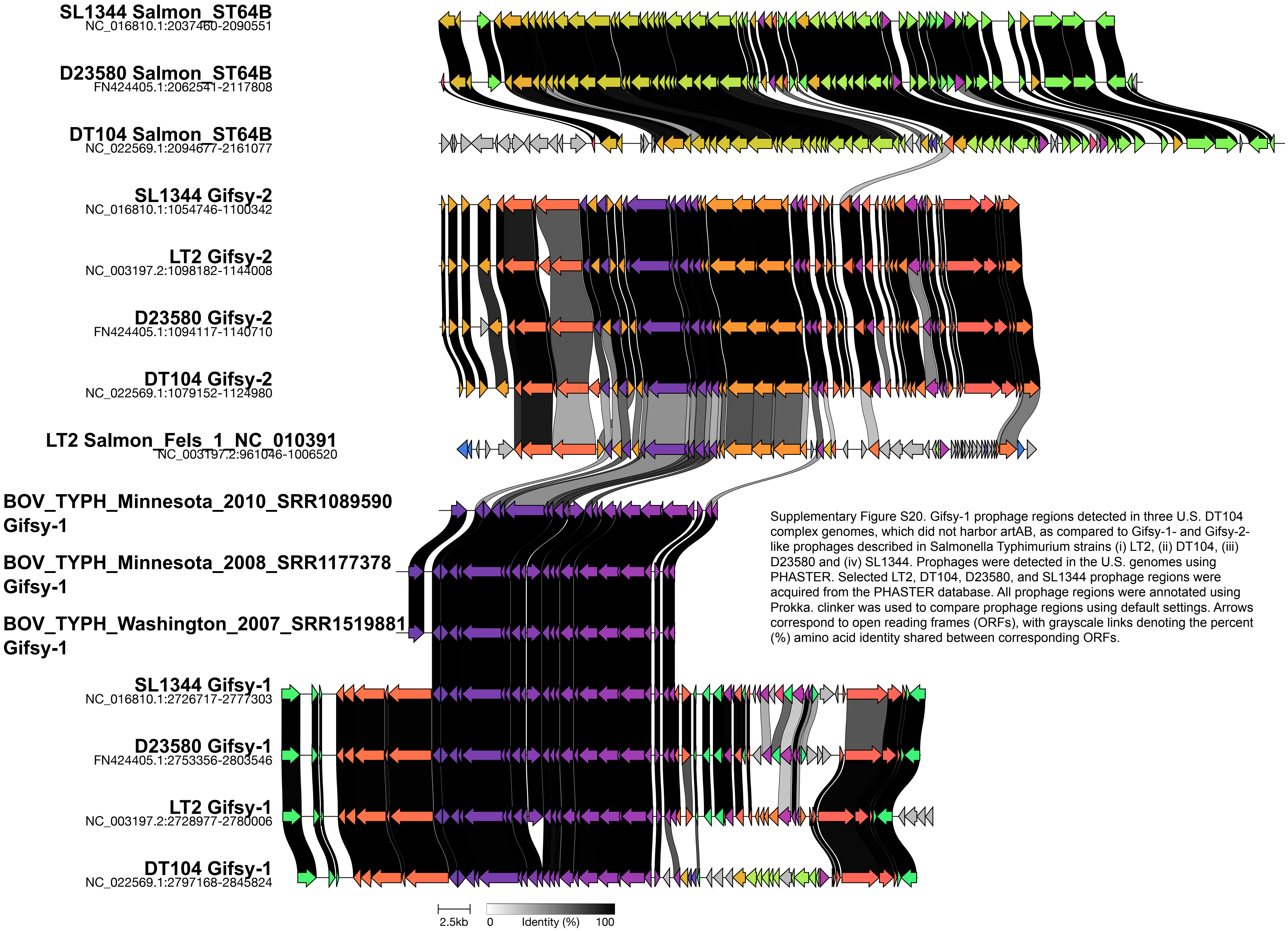


Supplementary Figure S20. Gifsy-1 prophage regions detected in three U.S. DT104 complex genomes, which did not harbor artAB, as compared to Gifsy-1- and Gifsy-2-like prophages described in Salmonella Typhimurium strains (i) LT2, (ii) DT104, (iii) D23580 and (iv) SL1344. Prophages were detected in the U.S. genomes using PHASTER. Selected LT2, DT104, D23580, and SL1344 prophage regions were acquired from the PHASTER database. All prophage regions were annotated using Prokka. clinker was used to compare prophage regions using default settings. Arrows correspond to open reading frames (ORFs), with grayscale links denoting the percent (%) amino acid identity shared between corresponding ORFs.
