## Supplementary Figure S21 for "A multidrug-resistant *Salmonella enterica* Typhimurium DT104 complex lineage circulating among humans and cattle in the United States lost the ability to produce pertussis-like toxin ArtAB"

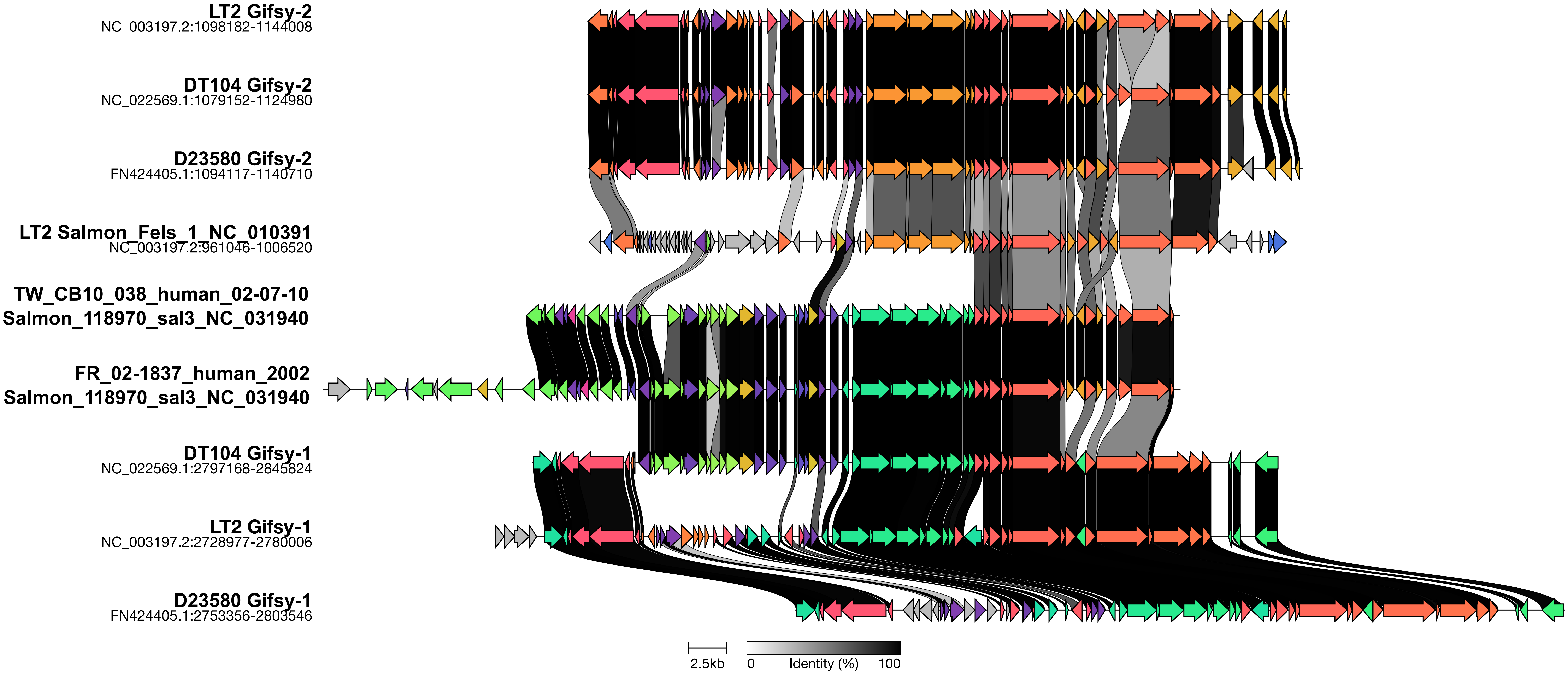


Supplementary Figure S21. artAB-harboring prophages detected in DT104 complex genomes, which were not labeled as “Gifsy-1” by PHASTER, as compared to Gifsy-1- and Gifsy-2-like prophages described in Salmonella Typhimurium strains (i) LT2, (ii) DT104, and (iii) D23580. Selected LT2, DT104, and D23580 prophage regions were acquired from the PHASTER database. All prophage regions were annotated using Prokka. clinker was used to compare prophage regions using default settings. Arrows correspond to open reading frames (ORFs), with grayscale links denoting the percent (%) amino acid identity shared between corresponding ORFs.
