## Supplementary Text for "A multidrug-resistant *Salmonella enterica* Typhimurium DT104 complex lineage circulating among humans and cattle in the United States lost the ability to produce pertussis-like toxin ArtAB"

for

by

Laura M. Carroll, Nicolo Piacenza, Rachel A. Cheng, Martin Wiedmann, Claudia Guldimann

### SUPPLEMENTARY METHODS

**Acquisition of U.S. human- and bovine-associated DT104 complex genomic data and metadata.** In a previous study of human- and bovine-associated *S.* Typhimurium from New York State, we identified three closely related, *artAB*-negative DT104 complex genomes from both humans and cattle (out of 14 total DT104 complex genomes from humans and cattle in New York State) [1]. Thus, as a first evaluation of *artAB* presence and absence in the DT104 complex, we compiled a set of DT104 complex genomes from humans and cattle across the U.S. (referred to hereafter as "Dataset 1 [U.S. bovine and human data]"; Figure 1B and Supplementary Figures S2 and S3).

To construct Dataset 1 (U.S. bovine and human data), we first collected genomic data derived from 14 human- and bovine-associated DT104 complex isolates from New York State, which we had sequenced in a previous study (members of the *S.* Typhimurium Lineage III cluster described in Supplementary Figures S2 and S5 of Carroll, et al.) [1]. We then aggregated these 14 New York State genomes with 223 human- and bovine-associated DT104 complex genomes from across the U.S., as described previously [1]. Briefly, paired-end Illumina short reads associated with 223 *S.* Typhimurium genomes meeting the following criteria were downloaded via Enterobase (accessed November 29, 2018) [2, 3] and the Sequence Read Archive (SRA) Toolkit version 2.9.3 [4, 5]: (i) genomes were serotyped as *S.* Typhimurium *in silico* using the implementation of SISTR [6] in Enterobase; (ii) the country of isolation was the United States; (iii) the isolation source was reported as either "Human" or "Bovine" in the "Source Niche" and "Source Type" fields in Enterobase, respectively; (iv) genomes had an isolation year reported in Enterobase; (v) using RhierBAPS [7], genomes were assigned to the DT104 complex, a well-supported cluster within the larger bovine- and human-associated U.S. *S.*

ABRicate version 0.8 [18] was used to detect antimicrobial resistance (AMR) genes, plasmid replicons, and virulence factors in each assembled DT104 genome using NCBI's National Database of Antibiotic Resistant Organisms (NDARO) [19], the PlasmidFinder database [20], and the Virulence Factor Database (VFDB) [21], respectively, using minimum nucleotide identity and coverage thresholds of 75 and 50%, respectively (all databases accessed December 10, 2020; Supplementary Table S3). The aforementioned ABRicate analyses were repeated, using a minimum coverage threshold of 0% (e.g., to confirm that virulence factors discussed in the manuscript were absent from genomes in which they were not initially detected).

Each assembled genome was additionally queried for the presence of selected virulence factors, which have previously been associated with prophage in *Salmonella* [22]: (i) *artAB* (NCBI Nucleotide Accession AB104436.1), (ii) *gogA* (European Nucleotide Archive [ENA] Accession EAA7850902.1), (iii) *gtgA* (ENA Accession PVI70081.1), and (iv) *gipA* (ENA Accession CAI93790.1). Assembled genomes were queried for selected virulence factors using the command-line implementation of nucleotide BLAST (blastn) version 2.11.0 [23], using default settings plus a minimum coverage threshold of 40% (Supplementary Table S3). To confirm that the aforementioned genes were absent from genomes in which they were not initially detected, all genomes were queried again (i) as described above, with the coverage threshold lowered to 0%; and (ii) using translated nucleotide BLAST (tblastx; Supplementary Tables S4 and S5).

ARIBA version 2.14.6 [24] was used to further confirm *artAB* and *gogB* presence/absence in all genomes with associated paired-end Illumina reads (*n* = 219 U.S. human- and bovine-associated DT104 complex genomes from Dataset 1 [U.S. bovine and human data] with paired-end Illumina reads; Figure 1B, Supplementary Figures S2 and S3, and Supplementary Tables S1 and S2). Briefly, the ARIBA database was constructed using the ARIBA "prepareref" command (default settings, except "--all_coding" set to "yes" and the "--no_cdhit" option included), and ARIBA was run using ARIBA's "run" command with default settings. *artAB* presence/absence results obtained via ARIBA were identical to results obtained via blastn (Supplementary Table S6). For *gogB*, ARIBA results matched BLAST results for all but five genomes (*n* = 214 of 219 genomes, 97.7%; Supplementary Table S6). For the five discordant genomes, *gogB* was detected via blastn, but not with ARIBA. Upon manual inspection of these five genomes, (i) three genomes had *gogB* split on multiple contigs, and (ii) two genomes had an intact *gogB* detected at high identity (>99.9% nucleotide identity) and 100% coverage via blastn. blastn results are thus reported in the main manuscript (Supplementary Tables S3-S5), with ARIBA results available in Supplementary Table S6.

**Variant calling and maximum likelihood phylogeny construction within Dataset 1 (U.S. bovine and human data).** Core single nucleotide polymorphisms (SNPs) were identified among all 230 genomes within Dataset 1 (U.S. bovine and human data), using the default pipeline implemented in Snippy version 4.6.0 [25] and the following dependencies: BWA version 0.7.17-r1188 [26, 27], Minimap2 version 2.23-r1111 [28], SAMtools version 1.14 [29], BEDtools version 2.30.0 [30, 31], BCFtools version 1.14 [32], FreeBayes version 1.3.2-dirty [33], vcflib version 1.0.0-rc0-349-g45c6-dirty [34], vt version 0.5 [35], SnpEff version 5.0e [36], samclip version 0.4.0 [37], seqtk version 1.3-r106 [38], and snp-sites version 2.5.1 [39]. For the 219 Dataset 1 (U.S. bovine and human data) genomes with paired-end Illumina reads available, the trimmed Illumina paired-end reads associated with each genome were used as input; for the remaining 11 genomes, the assembled contigs were used as input (see section "Acquisition of U.S. human- and bovine-associated DT104 complex genomic data and metadata" above; Supplementary Tables S1 and S2). The closed DT104 chromosome (NCBI Nucleotide accession NC_022569.1) was used as a reference. Core SNPs identified in regions of the DT104 chromosome predicted to belong to phage were masked (see section "*In silico* detection of prophage, antimicrobial resistance genes, plasmid replicons, and virulence factors" above). Gubbins version 2.4.1 [40] was used to identify and remove recombination events in all genomes using default settings, and snp-sites was used to query the resulting recombination-free alignment for core SNPs (i.e., using the "-c" option).

A maximum likelihood (ML) phylogeny was constructed with IQ-TREE version 1.5.4 [41], using (i) the resulting core SNPs as input, (ii) the optimal nucleotide substitution model determined using Bayesian information criteria (BIC) values produced with ModelFinder [42] (i.e., the K3Pu+I model) [43], (iii) an ascertainment bias correction to account for the use of solely variant sites (corresponding to constant sites identified relative to the DT104 reference chromosome; "-fconst 1092869,1195194,1193287,1094079"), and (iv) 1,000 replicates of the ultrafast bootstrap approximation [44, 45].

TempEst version 1.5.3 [46] was used to assess the temporal structure of the resulting unrooted ML phylogeny, using the best-fitting root and the $R^2$ function ($R^2 = 0.33$, slope = $3.05\times10^{-7}$ substitutions/site/year, X-intercept = 1988.1). The unrooted ML phylogeny was additionally rooted and time scaled using LSD2 version 1.4.2.2 [47] and the following parameters: (i) tip dates corresponding to the year of isolation associated with each genome; (ii) an estimated substitution rate; (iii) constrained mode ("-c"), with the root estimated using constraints on all branches ("-r as"); (iv) variances calculated using input branch lengths ("-v 1"); (v) 1,000 samples for calculating confidence intervals for estimated dates ("-f 1000"); (vi) a sequence length of 4,500,000. The resulting rooted, time-scaled ML phylogeny was viewed using FigTree version 1.4.4 [48] (Supplementary Data).

**Dataset 1 (U.S. bovine and human data) Bayesian time-scaled phylogeny construction.** In addition to constructing a time-scaled ML phylogeny (see section "Variant calling and maximum likelihood phylogeny construction within Dataset 1 [U.S. bovine and human data]" above), a Bayesian approach was additionally employed to construct time-scaled phylogenies for Dataset 1 (U.S. bovine and human data). Due to an overrepresentation of strains reportedly isolated in 2007 from bovine sources in Washington State within Dataset 1 (U.S. bovine and human data) (*n*

= 94 of 230 Dataset 1 [U.S. bovine and human data] genomes, 40.9%; Figure 2, Supplementary Figure S4, and Supplementary Table S1), all aforementioned SNP calling and ML phylogeny construction steps were repeated among Dataset 1 (U.S. bovine and human data) genome sets downsampled to (i) 25, (ii) 10, and (iii) 5 randomly selected bovine genomes collected in Washington State in 2007 (to minimize the risk of parameter estimate biases during Bayesian phylogeny construction, $n$ = 161, 146, and 141 total genomes in each downsampled genome set, respectively, out of 230 total Dataset 1 [U.S. bovine and human data] genomes; see section "Variant calling and maximum likelihood phylogeny construction within Dataset 1 [U.S. bovine and human data]" above, Supplementary Data) [49]. All resulting ML phylogenies were time-scaled using TempEst and LSD2 as described above (see section "Variant calling and maximum likelihood phylogeny construction within Dataset 1 [U.S. bovine and human data]" above; Supplementary Table S7).

For each of the three downsampled Dataset 1 (U.S. bovine and human data) genome sets, BEAST2 version 2.5.1 [50, 51] was used to construct a tip-dated phylogeny, using core SNPs detected among the genomes within the respective downsampled data set as input (see section "Variant calling and maximum likelihood phylogeny construction within Dataset 1 [U.S. bovine and human data]" above). For all three downsampled genome sets, an initial clock rate of $2.79 \times 10^{-7}$ substitutions/site/year [13] was used, along with an ascertainment bias correction to account for the use of solely variant sites [52]. bmodeltest [53] was used to infer a substitution model using Bayesian model averaging, with transitions and transversions split. A relaxed lognormal molecular clock [54] and a coalescent Bayesian skyline population model [55] were used, as these models have been selected as the optimal clock/population model combination for DT104 previously [13]. A log-normal distribution with a mean of $4.6 \times 10^{-7}$ and standard deviation of 1 (median of $2.79\times10^{-7}$) was used as the prior on the uncorrelated log-normal relaxed molecular clock mean rate parameter (ucld.mean; Supplementary Data).

For each of the three downsampled Dataset 1 (U.S. bovine and human data) genome sets, five independent BEAST2 runs (i.e., BEAST2 runs with different random seeds) were performed, using chain lengths of at least 100 million generations, sampling every 10 thousand generations. For each downsampled data set, LogCombiner-2 was used to aggregate the resulting log and tree files with 10% of the states treated as burn-in, and TreeAnnotator-2 was used to produce a maximum clade credibility (MCC) tree using Common Ancestor node heights (Supplementary Data). The resulting phylogenies were displayed and annotated using R version 4.1.2 [56] and the following packages: ggplot2 version 3.3.5 [57], ggtree version 3.2.1 [58, 59], phylobase version 0.8.10 [60], and treeio version 1.18.1 [61].

All three downsampled Dataset 1 (U.S. bovine and human data) genome sets resulted in similar BEAST2 parameter estimates (Supplementary Figure S5, Supplementary Table S8, and Supplementary Data). The final Bayesian time-scaled phylogeny and associated parameter estimates reported in the main manuscript correspond to those obtained using the Dataset 1 (U.S. bovine and human data) genome set, which was downsampled to 10 randomly selected bovine genomes collected in Washington State in 2007 ($n$ = 146 genomes, Figures 3 and 4). Results are available for the Dataset 1 (U.S. bovine and human data) genome sets downsampled to 25 and 5 bovine genomes collected in Washington State in 2007 (Supplementary Figures S6-S9, Supplementary Table S8, and Supplementary Data).

***artAB* ancestral state reconstruction for Dataset 1 (U.S. bovine and human data).** To estimate ancestral character states of internal nodes in the Dataset 1 (U.S. bovine and human data) phylogeny as they related to *artAB* presence/absence (i.e., whether a node in the tree represented an ancestor that was more likely to be *artAB*-positive or *artAB*-negative), the presence or absence of *artAB* within each genome was treated as a binary state (see section "*In silico* detection of prophage, antimicrobial resistance genes, plasmid replicons, and virulence factors" above). Three separate *artAB* ancestral state reconstruction runs were performed, each using one of the three BEAST2 time-scaled Bayesian Dataset 1 (U.S. bovine and human data) phylogenies as input ($n$ = 161, 146, and 141 total genomes in each downsampled Dataset 1 [U.S. bovine and human data] genome set; see section "Dataset 1 [U.S. bovine and human data] Bayesian time-scaled phylogeny construction" above).

Stochastic character maps were simulated on each phylogeny using the make.simmap function in the phytools version 1.0-1 R package [62] and the all-rates-different (ARD) model in the ape version 5.6-1 package [63, 64]. For each phylogeny, either (i) equal root node prior probabilities for *artAB*-positive and *artAB*-negative states (i.e., $P$(*artAB* present) = $P$(*artAB* absent) = 0.5), or (ii) estimated root node prior probabilities for *artAB*-positive and *artAB*-negative states obtained using the make.simmap function were used. For each root node prior/phylogeny combination (six total combinations of two root node priors and three Dataset 1 [U.S. bovine and human data] phylogenies), an empirical Bayes approach was used, in which a continuous-time reversible Markov model was fitted, followed by 10,000 simulations of stochastic character histories using the fitted model and tree tip states. The resulting phylogenies were plotted using the densityMap function in the phytools R package. For Dataset 1 (U.S. bovine and human data), the final ancestral state results reported in the main manuscript correspond to those obtained using the Dataset 1 (U.S. bovine and human data) genome set, which was downsampled to 10 randomly selected bovine genomes collected in Washington State in 2007 ($n$ = 146 genomes, Figure 3). Results are available for the Dataset 1 (U.S. bovine and human data) genome sets downsampled to 25 and 5 bovine genomes collected in Washington State in 2007 (Supplementary Figures S10-S12 and Supplementary Data).

**Pan-genome characterization of Dataset 1 (U.S. bovine and human data).** Prokka version 1.13.3 [16] was used to annotate all 230 genomes within Dataset 1 (U.S. bovine and human data), using the "Bacteria" database and default settings (Supplementary Tables S1 and S2). GFF files produced by Prokka were supplied as input to Panaroo version 1.2.7 [65], which was used to identify core- and pan-genome orthologous gene clusters among the 230 Dataset 1 (U.S. bovine and human data) genomes, with the following parameters: (i) "strict" mode ("--clean-mode strict"); (ii) MAFFT as the sequence aligner ("--aligner mafft") [66, 67]; (iii) a core genome threshold of 98% (i.e., genes present in at least 98% of genomes were considered to be core genes; "--core_threshold 0.98"); (iv) a protein family sequence identity threshold of 70% ("-f 0.7", the default). The LSD2 time-scaled ML phylogeny for Dataset 1 (U.S. bovine and human data) (see section "Variant calling and maximum likelihood phylogeny construction within Dataset 1 [U.S. bovine and human data]" above) was supplied as input to Panaroo's "panaroo-img" and "panaroo-fmg" commands, which were used to estimate the pan-genome size under the Infinitely Many Genes (IMG) [68, 69] and Finite Many Genes (FMG) models (with 100 bootstrap replicates) [70], respectively (Supplementary Figure S13).

Reference pan-genome coding sequences (CDS) identified by Panaroo underwent functional annotation using the eggNOG-mapper version 2 webserver (http://eggnog-mapper.embl.de/; accessed July 24, 2022) using default settings [71, 72]. The "table" function in R was used to identify genes associated with (i) prophage Gifsy-1 presence/absence (Supplementary Table S9) and (ii) clade membership (Supplementary Table S10); the "fisher.test" function in R's stats package was used to conduct two-sided Fisher's exact tests, and the “p.adjust” function was used to control the false discovery rate (i.e., p.adjust method = “fdr”) [73].

**Genome-wide identification of host-associated orthologous gene clusters for Dataset 1 (U.S. bovine and human data).** The treeWAS version 1.0 R package [74] was used to identify potential orthologous gene cluster-host associations among the 230 human- and bovine-associated U.S. DT104 complex genomes in Dataset 1 (U.S. bovine and human data) (i.e., whether an orthologous gene cluster identified with Panaroo was human- or bovine-associated while accounting for population structure). The following treeWAS parameters were used: (i) the isolation source was treated as a discrete phenotype (i.e., a vector of “human” or “bovine”, supplied to the “treeWAS” function’s “phen” argument; phen.type = “discrete”); (ii) unique gene presence/absence profiles of genes detected in ≥10 and ≤220 of 230 total Dataset 1 (U.S. bovine and human data) genomes, treated as the genotypes to test (supplied to the treeWAS function’s “snps” argument); (iii) the time-scaled ML phylogeny constructed using LSD2 was supplied as input to the “treeWAS” function’s “tree” argument (see section “Variant calling and maximum likelihood phylogeny construction within Dataset 1 [U.S. bovine and human data]” above); (iv) the number of simulated loci for estimating the null distribution was set to five million (i.e., n.snps.sim = 5000000); (v) ancestral state reconstruction performed using ML methods (i.e., snps.reconstruction = “ML”, snps.sim.reconstruction = “ML”, and phen.reconstruction = “ML”); (vi) a *P*-value significance threshold of 0.1, after controlling the FDR (p.value.correct = “fdr”). The analysis was re-run, using parsimony approaches in place of ML approaches for ancestral state reconstruction. Regardless of approach, no orthologous gene clusters were found to be significantly associated with isolation source via any of the treeWAS association tests (FDR-corrected *P*-value > 0.1).

**Acquisition of global DT104 complex genomic data and metadata.** To compare the 230 U.S. human- and bovine-associated DT104 complex genomes in Dataset 1 (U.S. bovine and human data) to a larger set of DT104 complex genomes from numerous sources worldwide, genomic data associated with the following studies were downloaded via Enterobase: (i) Illumina reads associated with 243 bovine- and human-associated DT104 isolates from a study of between-host transmission within Scotland [75] (referred to hereafter as "Dataset 2 [Scottish bovine and human data]"); genomes were pre-processed and assembled as described above (see section "Acquisition of U.S. human- and bovine-associated DT104 complex genomic data and metadata" above); (ii) assembled genomes associated with 290 DT104 isolates from a variety of sources and countries from a study describing the global spread of DT104 [13] (referred to hereafter as "Dataset 3 [multi-source data]"; eleven of the 290 genomes were isolated from cattle and humans in the U.S. and thus had also been included in Dataset 1 [U.S. bovine and human data ], Figure 1B, Supplementary Figures S2 and S3, and Supplementary Table S1).

The following datasets were aggregated to create a final set of 752 DT104 complex genomes derived from multiple different countries and isolation sources, which was used in subsequent steps (referred to hereafter as "Dataset 4 [combined global dataset]"; Figure 1B, Supplementary Figures S2 and S3, and Supplementary Tables S1 and S2): (i) Dataset 1 (U.S. bovine and human data) ($n$ = 230 DT104 complex genomes), (ii) Dataset 2 (Scottish bovine and human data) ($n$ = 243 DT104 genomes), and (iii) Dataset 3 (multi-source data) ($n$ = 290 DT104 genomes, including 11 genomes that were part of Dataset 1 [U.S. bovine and human data]). QUAST version 4.5 was used to assess the quality of all 752 genomes in Dataset 4 (combined global dataset) (Supplementary Tables S1 and S2). Prophage, AMR genes, plasmid replicons, and virulence factors were detected in all 752 Dataset 4 (combined global dataset) genomes as described above (see section "*In silico* detection of prophage, antimicrobial resistance genes, plasmid replicons, and virulence factors" above; Supplementary Tables S3-S5).

**Variant calling and maximum likelihood phylogeny construction within Dataset 4 (combined global dataset).** To identify core SNPs present in all 752 DT104 complex genomes within Dataset 4 (combined global dataset), Parsnp and HarvestTools version 1.2 [76] were used, as Parsnp easily scales to large data sets (Supplementary Tables S1 and S2) [76]. Assembled genomes were used as input for Parsnp, along with the closed DT104 chromosome as a reference (NCBI Nucleotide accession NC_022569.1) and Parsnp's implementation of PhiPack [77] to filter recombination.

Core SNPs detected among all 752 assembled genomes within Dataset 4 (combined global dataset) were supplied as input to IQ-TREE version 1.5.4, which was used to construct a ML phylogeny as described above (the corresponding ascertainment bias correction here was "-fconst 1181208,1285673,1280769,1179580"; see section "Variant calling and maximum likelihood phylogeny construction within Dataset 1 [U.S. bovine and human data]" above). The resulting ML phylogeny was rooted and time-scaled using LSD2 as described above (see section "Variant calling and maximum likelihood phylogeny construction within Dataset 1 [U.S. bovine and human data]" above; Supplementary Data). A range of 1900-2017 was supplied for four genomes, which were part of Dataset 3 (multi-source data), but did not have a reported year of isolation. The resulting LSD2 time-scaled ML phylogeny was annotated using the Interactive Tree of Life (iTOL) version 6 webserver (https://itol.embl.de/, accessed March 7, 2022; Figure 5, Supplementary Figure S14, Supplementary Data) [78]. The LSD2 time-scaled ML phylogeny for Dataset 4 (combined global dataset) was further used for *artAB* presence/absence ancestral state reconstruction as described above (see section "*artAB* ancestral state reconstruction for Dataset 1 [U.S. bovine and human data]" above; Supplementary Figure S15).

**Pan-genome characterization of Dataset 4 (combined global dataset).** Pan-genome analyses were carried out for Dataset 4 (combined global dataset) as described above (see section "Pan-genome characterization of Dataset 1 [U.S. bovine and human data]" above; Supplementary Tables S1 and S2). Briefly, Prokka was used to annotate all 752 genomes within Dataset 4 (combined global dataset). GFF files produced by Prokka were supplied as input to Panaroo, which was used to identify core- and pan-genome orthologous gene clusters among all 752 Dataset 4 (combined global dataset) genomes. The pan-genome size for Dataset 4 (combined global dataset) was estimated using Panaroo's "panaroo-img" and "panaroo-fmg" commands, using the LSD2 time-scaled ML phylogeny for Dataset 4 (combined global dataset) as input (Supplementary Figure S13). Reference pan-genome CDS identified by Panaroo underwent functional annotation using eggNOG-mapper.

**Strain selection for phenotypic stress assays.** Phenotypic stress assays (discussed in detail in the sections below) were used to compare (i) bovine- and human-associated, Gifsy-1/*artAB*/*gogB*-positive U.S. DT104 complex strains to (ii) bovine- and human-associated, Gifsy-1/*artAB*/*gogB*-negative U.S. DT104 complex strains. Thus, the genomes of 13 bovine- and human-associated DT104 complex strains from New York State [1], which were available to us in the Cornell University Food Safety Laboratory (CUFSL) culture collection [79], were characterized further (Supplementary Figures S2 and S3 and Supplementary Table S2).

To identify a set of closely related, Gifsy-1/*artAB*/*gogB*-positive and -negative genomes for experimental characterization, Parsnp and HarvestTools version 1.2 [76] were used to identify core SNPs among all 13 genomes of New York State DT104 complex strains available in the CUFSL culture collection (Supplementary Figures S2 and S3 and Supplementary Table S2). Assembled genomes were supplied as input to Parsnp, along with the closed DT104 chromosome as a reference (NCBI Nucleotide accession NC_022569.1) and Parsnp's implementation of PhiPack [77] to remove recombination. IQ-TREE version 1.5.4 was used to construct a ML phylogeny, using (i) the resulting core SNPs as input, (ii) an ascertainment bias correction, based on the GC content of the DT104 reference chromosome ("-fconst 1182070,1287912,1283169,1180480"), (iii) the optimal nucleotide substitution model ("-m MFP"), selected using ModelFinder (i.e., the TIM+I model), and (iv) 1,000 replicates of the ultrafast bootstrap approximation ("-bb 1000").

Further steps were taken to ensure that the selected DT104 complex strains were as similar as possible in terms of their pan-genome composition. Briefly, Prokka version 1.13 was used to annotate each genome (using the "Bacteria" database, plus default settings). The resulting GFF files were supplied to Roary version 3.13.0 [80], which was used to identify orthologous gene clusters among the 13 DT104 complex genomes available for experimental characterization (using default thresholds, e.g., 95% protein BLAST [blastp] identity; Supplementary Table S2).

The New York State DT104 complex genomes differed little in terms of their core and pan-genome compositions (Supplementary Figure S16 and Supplementary Table S2). A total of 336 core SNPs were identified among the 13 DT104 complex genomes; pairwise core SNP distances between all 13 genomes ranged from 12-113 core SNPs (median and mean of 85 and 80.8 core SNPs, respectively, calculated using the "dist.gene" function in the ape R package). Based on gene presence/absence of pan-genome elements identified via Roary, the Jaccard distance between all 13 genomes ranged from 0.0036-0.0820 (median and mean of 0.0359 and 0.0380, respectively; calculated in R using the "vegdist" function in vegan version 2.5-7) [81].

Considering both (i) core- and pan-genome similarities between all 13 available New York State DT104 complex genomes, as well as (ii) Gifsy-1/*artAB*/*gogB* presence and absence, we selected six closely related, New York State DT104 complex strains to undergo phenotypic characterization (i.e., three Gifsy-1/*artAB*/*gogB*-positive strains, and three Gifsy-1/*artAB*/*gogB*-negative strains; Supplementary Table S11). Briefly, all available Gifsy-1/*artAB*/*gogB*-negative strains in the CUFSL culture collection were selected to undergo phenotypic testing ($n$ = 3, two human isolates and one bovine isolate; Supplementary Table S11); all three strains were members of the U.S. *artAB*-negative major clade (discussed in detail in the "Results" section below). Considering both core- and pan-genome distances relative to all three available Gifsy-1/*artAB*/*gogB*-negative strains, three Gifsy-1/*artAB*/*gogB*-positive DT104 complex strains were additionally selected to undergo phenotypic testing (one from human and two from bovine sources; Figure 5, Supplementary Figure S16, and Supplementary Table S11). The three selected Gifsy-1/*artAB*/*gogB*-positive DT104 complex strains differed from the three available Gifsy-1/*artAB*/*gogB*-negative strains by (i) 64-83 (HUM_TYPH_NY_04_S5_0370), 74-93 (BOV_TYPH_NY_99_A4_0023), and 65-84 (BOV_TYPH_NY_99_S3_0910) core SNPs and (ii) Jaccard distances (based on pan-genome element presence/absence) of 0.0148-0.0610 (HUM_TYPH_NY_04_S5_0370), 0.0174-0.0622 (BOV_TYPH_NY_99_A4_0023) and 0.0163-0.0620 (BOV_TYPH_NY_99_S3_0910; Figure 5, Supplementary Figure S16, and Supplementary Table S11).
